# Mitochondrial fission, integrity and completion of mitophagy require separable functions of Vps13D in *Drosophila* neurons

**DOI:** 10.1101/2020.01.21.914523

**Authors:** Ryan Insolera, Péter Lőrincz, Alec J Wishnie, Gábor Juhász, Catherine A Collins

**Affiliations:** Molecular, Cellular and Developmental Biology Department, University of Michigan, Ann Arbor, MI; Department of Anatomy, Cell and Developmental Biology, Eötvös Loránd University, Budapest, H-1117 Hungary; Premium Postdoctoral Research Program, Hungarian Academy of Sciences, Budapest, Hungary; Institute of Genetics, Biological Research Centre, Szeged, H-6726 Hungary

## Abstract

A healthy population of mitochondria, maintained by proper fission, fusion, and degradation, is critical for the long-term survival and function of neurons. Here, our discovery of mitophagy intermediates in fission-impaired *Drosophila* neurons brings new perspective into the relationship between mitochondrial fission and mitophagy. Neurons lacking either the ataxia disease gene Vps13D or the dynamin related protein Drp1 contain enlarged mitochondria that are engaged with autophagy machinery and also lack matrix components due to rupture. Reporter assays combined with genetic studies imply that mitophagy both initiates and is completed in Drp1 impaired neurons, but fails to complete in Vps13D impaired neurons, which accumulate compromised mitochondria within stalled mito-phagophores. Our findings imply that in fission-defective neurons, mitophagy becomes induced, and that the lipid channel containing protein Vps13D has separable functions in mitochondrial fission and phagophore elongation.

## Introduction

Neurons are among the most sensitive cell types to mitochondrial perturbation, and are heavily reliant upon a proper balance of mitochondrial biogenesis and degradation, as well as fission and fusion dynamics (Misgeld and Schwarz 2017). Mutations that disrupt mitochondrial fission and fusion machinery lead to severe neurological dysfunction in humans and animal models (Burté et al. 2015; Detmer et al. 2008; Davies et al. 2007). Likewise, quality control mechanisms, including the autophagic clearance of damaged mitochondria, known as mitophagy (Pickles, Vigié, and Youle 2018), play important protective roles in neurons; impaired mitophagy is thought to contribute to pathology of multiple neurodegenerative diseases (Markaki and Tavernarakis 2020).

Research in the past decade has uncovered numerous cellular components of mitophagy machinery, primarily through studies that follow toxin-induced damage to the entire mitochondrial population (D. Narendra et al. 2008; Geisler et al. 2010; D. P. Narendra et al. 2010; Cai et al. 2012). However, neurons strongly require mitochondria, so are unlikely to undergo widespread clearance of mitochondria or survive such harsh insults (Whitworth and Pallanck 2017; Cummins and Götz 2018). Instead, neurons are expected to selectively degrade only damaged mitochondria, however molecular tools to study this type of mitophagy in neurons have been limited.

The development of specialized acid-sensitive fluorescent reporters have opened opportunities to monitor mitophagy *in vivo* (McWilliams et al. 2016; Cao et al. 2017; Lee et al. 2018; Sun et al. 2015), and have thus far been tested in conditions shown to alter *in vitro* toxin-induced mitophagy, with mixed results. Different reporters, mitoQC and mitoKeima, targeted to the outer mitochondrial membrane (OMM) or matrix, respectively (McWilliams et al. 2018; Lee et al. 2018; Cornelissen et al. 2018), yielded different interpretations of *in vivo* phenotypes. While some of these differences can be attributed to the use of different reporters and model organisms (Rodger, McWilliams, and Ganley 2018), the range of differences in existing studies emphasizes the remaining large gap in our understanding of mitophagy mechanisms in neurons.

Another potential discrepancy between mitophagy in cultured cells compared to neurons *in vivo* is the understood role of mitochondrial fission. Multiple studies (Twig et al. 2008; Tanaka et al. 2010; Frank et al. 2012), with one exception (Yamashita et al. 2016), have indicated that mitochondrial fission is required for the induction of mitophagy in cultured cells subjected to toxins (Burman et al. 2017; Twig et al. 2008; MacVicar and Lane 2014; Tanaka et al. 2010). Conditional knockout of the essential fission protein dynamin-related protein 1 (Drp1) in Purkinje cells of the mouse cerebellum results in the accumulation of autophagy components (ubiquitin, p62, and LC3) on mitochondria (Kageyama et al. 2012, 2014). These observations suggest that fission via Drp1 is not required for the initiation of mitophagy in neurons, however an understanding of Drp1’s role requires a better understanding of what these autophagy-targeted mitochondria, termed halted mitophagy intermediates (Yamada et al. 2019), represent. One possibility is that Drp1 loss leads to induction of mitophagy, which allows detection of the intermediates because of their frequency. An opposite possibility, which was inferred by these reports, is that Drp1 loss leads to a block in the progression of mitophagy, causing the accumulation of stalled intermediates in the cerebellum. Further *in vivo* studies are needed, with the acknowledgement that mechanisms that support the survival of neurons over the long life time of an animal may diverge considerably from mechanisms sufficient in cultured cells.

With powerful genetic and cell biological tools, *Drosophila melanogaster* is an ideal model organism to study mitochondrial quality control mechanisms in the nervous system. Our current study investigates the role and relationship of mitochondrial fission with mitophagy in the nervous system. Our starting point was to understand the neuronal function of the neurodegenerative disease-associated protein Vps13D (Vacuolar Protein Sorting protein 13D). The Vps13 protein family (Vps13A-D) has been characterized as phospholipid transporters that specifically localize to inter-organelle contact sites (Kumar et al. 2018; P. Li et al. 2020; Bean et al. 2018; Yeshaw et al. 2019; Park et al. 2016). While a specific cellular function has not yet been established for each of the Vps13 proteins, it is clear that they are all critical for neuronal health, as mutations in all family members are associated with neurological disorders in humans (Rzepnikowska et al. 2017; Seong et al. 2018; Gauthier et al. 2018).

In 2018 our collaborators and others identified *VPS13D* as a cause of familial ataxia (Seong et al. 2018), developmental movement disorders (Seong et al. 2018; Gauthier et al. 2018) and spastic paraplegia (Koh et al. 2020). We found that Vps13D depleted neurons accumulate severely enlarged mitochondria which fail to be trafficked to distal axons (Seong et al. 2018). Indeed, loss of Vps13D function in many different cell types leads to severely enlarged mitochondria, including *Drosophila* intestinal cells, HeLa cells (Anding et al. 2018), and cultured fibroblasts from human ataxia patients containing point mutations in the *VPS13D* gene (Seong et al. 2018). These findings suggest a conserved role for Vps13D in mitochondrial dynamics.

However, in neurons we noticed additional phenotypes associated with loss of Vps13D: here we report the accumulation of objects that appear to be stalled intermediates that have initiated but fail to complete mitophagy, and are also ruptured such that matrix components are lost. The loss of matrix makes them less obvious to detect by traditional means of mitochondrially targeted reporters, but having described these objects, we were then able to detect similar objects in neurons mutant or depleted for Drp1. In comparing mitophagy in the fission-deficient conditions of Vps13D and Drp1 depletion, we found that: (1) neurons deficient in their capacity for mitochondrial fission can still undergo functional mitophagy; (2) An additional function is revealed for Vps13D in phagophore elongation, and this role is independent and separable from a role in mitochondrial fission; and (3) impairment in mitochondrial fission, via loss of Drp1 and/or Vps13D function, is now established to be a *bona fide* stress that leads to an induction of mitophagy of individual mitochondria in neurons *in vivo*.

## Results

### Neurons lacking Vps13D contain rupturing mitochondria engaged with a phagophore

Consistent with our previous work (Seong et al. 2018), and previous observations in intestinal cells (Anding et al. 2018), RNAi depletion of Vps13D in larval motoneurons led to severely enlarged neuronal mitochondria, as visualized by Gal4-driven expression of matrix-targeted GFP (mitoGFP) (Rizzuto et al. 1995; Pilling et al. 2006) (**Figure 1A**). We also noticed large, round polyubiquitin (PolyUb) positive objects that were of similar size to the enlarged mitochondria, but lacked mitoGFP (**Figure 1A**). The PolyUb+ objects were also strongly positive for Ref(2)p the Drosophila homolog of mammalian p62 (**Figure 1A**).

**Figure 1:**
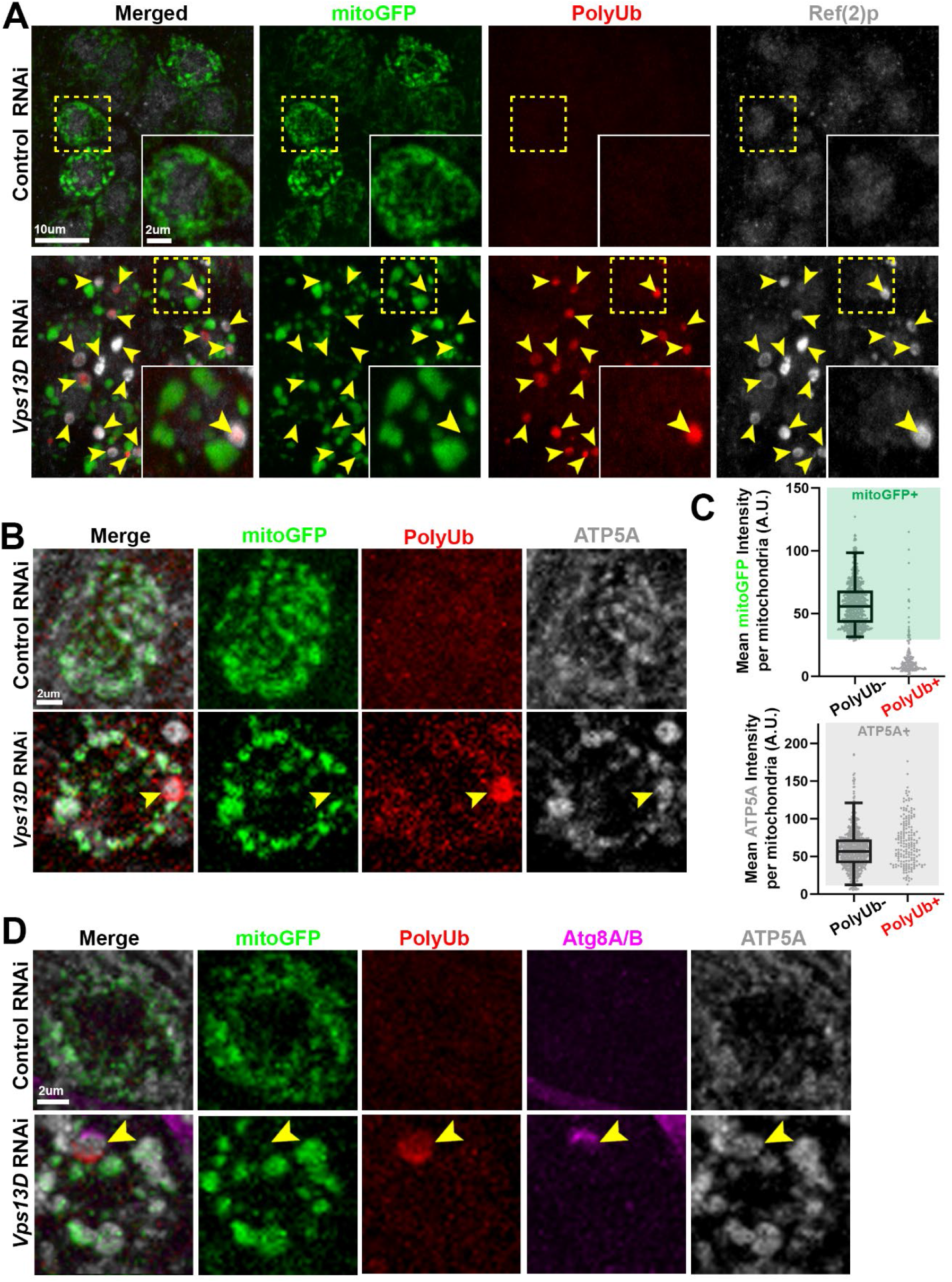
Neurons depleted for Vps13D contain mitophagy intermediates lacking matrix. Representative images of motoneurons in the larval VNC which co-express UAS-mitoGFP (green) with UAS-luciferase RNAi (control) (BL# 31603) versus UAS-*Vps13D* RNAi (BL# 38320), via the D42-Gal4 driver. **A)** VNC tissue was stained for polyubiquitin (PolyUb, FK1, red) and Ref(2)p (p62 homolog, greyscale). Dashed yellow box outlines a single Gal4-expressing neuronal cell body that is shown in high magnification in the inset in the bottom right corner. Yellow arrowheads indicate examples of PolyUb+ objects (PolyUb+/Ref(2)p+/mitoGFP-). *Vps13D* RNAi expressing neurons contained on average 1.88 PolyUb+ objects per cell body (n=239 neurons from 5 ventral nerve cords (VNCs)), while such objects were never observed in neurons expressing control RNAi. 98.8% of PolyUb+ objects were Ref(2)p+, n=485 PolyUb+ objects from 5 larval VNCs. **B)** Co-staining for mitoGFP (green), polyubiquitin (PolyUb, red) and endogenous IMM protein ATP5A (greyscale). The yellow arrowhead highlights an example PolyUb+ object, which contains ATP5A, but lacks mitoGFP. **C)** Distribution of mean intensities for mitoGFP (top) and ATP5A (bottom) from mitochondria in motoneurons. The right column shows objects selected based on positivity for PolyUb. The left column shows the population of conventional (PolyUb-) mitochondria (selected by similar criteria). Based on the criteria of >2.5% of the intensity distribution of the PolyUb- mitochondria population (shaded boxes), PolyUb+ objects were classified as + for mitoGFP and/or ATP5A if they were within the shaded box. n=809 mitochondria and 184 PolyUb+ objects from 5 larval VNCs. **D)** Co-staining for polyubiquitin (PolyUb, red), endogenous Atg8A/B (magenta), and endogenous IMM protein ATP5A (greyscale). Yellow arrowhead indicates an example of a mitophagy intermediate engaged with a phagophore but lacking mitochondrial matrix (PolyUb+/ATP5A+/Atg8+/mitoGFP-). 67.7% of the PolyUb+/ ATP5A+ objects in the neurons lacking Vps13D showed co-localization with Atg8A/B. n=425 PolyUb+/ATP5A+ objects from 5 larval VNCs.

We hypothesized that the PolyUb+/Ref(2)p+ objects are comprised of cellular substrates that are incompletely degraded in conditions of Vps13D depletion. To our surprise, we observed that PolyUb+ objects stained positive for the endogenous inner-mitochondrial membrane (IMM) protein ATP5A (**Figure 1B**). Of these enlarged PolyUb+ objects in the neuronal cell bodies, only 9.8% had mitoGFP intensities that fell within the range of the traditional neuronal mitochondria in this condition, which were PolyUb-, but contained clear mitoGFP and ATP5A staining (**Figure 1C**). However, all (100%) of these PolyUb+ objects had ATP5A intensities within the range of these traditional neuronal mitochondria. We observed similar Ref(2)p+/ATP5A+ objects in neurons of *Vps13D* mutant larvae (**Figure 1, Supplement 1A**), which died in early larval stages (Anding et al. 2018; Seong et al. 2018).

Since the mitoGFP marker is derived from an exogenously expressed transgene, we probed for additional mitochondrial components. The endogenous matrix enzyme pyruvate dehydrogenase E1ɑ (PDH) was also strongly reduced in the PolyUb+ objects (**Figure 1, Supplement 2A&B**). In addition, exogenous expression of an epitope tagged version of the full-length matrix enzyme isocitrate dehydrogenase 3β (UAS-Idh3b-HA) (Duncan, Kiefel, and Duncan 2017) labels mitochondria but failed to label the PolyUb+ objects (**Figure 1, Supplement 2C**). From these observations we therefore infer that the PolyUb+/Ref(2)p+/ATP5A+/mitoGFP- objects are derived from mitochondria that have somehow lost soluble matrix components.

The presence of the autophagy receptor Ref(2)p suggested that the PolyUb+/Ref(2)p+/ATP5A+/mitoGFP- objects may be in the process of mitophagy. We therefore probed for the presence of the core phagophore component Atg8, (homologous to LC3) (Gatica, Lahiri, and Klionsky 2018), using antibodies that recognize endogenous Atg8A/B (Barekat et al. 2016) (**Figure 1D**). We observed that 67.7% of the PolyUb+/ ATP5A+ objects in the neurons lacking Vps13D showed colocalization with Atg8A/B. Similar colocalization of ATP5A and Atg8A/B was observed in *Vps13D* mutant neurons (**Figure 1, Supplement 1B**), while no comparable structures were observed in control animals.

Since a large fraction of the PolyUb+/Ref(2)p+/ATP5A+/mitoGFP- objects associate with Atg8, and these objects are abundant in the ventral nerve cord (VNC) of larvae expressing *Vps13D* RNAi under the pan-neuronal elav-Gal4 driver, we expected that they would be detectable by electron microscopy (EM) based on the presence of the double membrane phagophore. We first observed the striking enlargement in mitochondrial morphology in Vps13D depleted neurons compared to VNCs expressing control RNAi (**Figure 2A**), which is similar to the mitochondrial phenotype in Vps13D depleted intestinal cells (Anding et al. 2018). The micrographs also revealed that some of the enlarged mitochondria were associated with a non-electron-dense double membrane consistent with a phagophore (designated as M_p_ in **Figure 2B**, with other examples in **Figure 2, Supplement 1A**). Mitochondria engaged with a phagophore were consistently more disorganized, and less electron dense than neighboring mitochondria in the same micrograph (**Figure 2, Supplement 1A**). Strikingly, in some instances we observed clear membrane rupture and loss of barrier between the outer-mitochondrial membrane (OMM), IMM, and the cytoplasm (**Figure 2C**, with other examples in **Figure 2, Supplement 1B**). While clear rupture was only observed in a subpopulation of the phagophore-associated mitochondria (3 out of 14), the micrographs capture only a single plane of a large three-dimensional mitochondrion, hence could miss rupture in a different location. Consistent with this notion, we observed additional mitochondria which resembled the phagophore-associated mitochondria in their more loosely packed and less electron-dense cristae (**Figure 2, Supplement 1C**). We hypothesize that the additional mitochondria in Vps13D depleted neurons that show reduced electron density may represent additional examples of the Atg8+ mitochondria observed by light microscopy.

**Figure 2:**
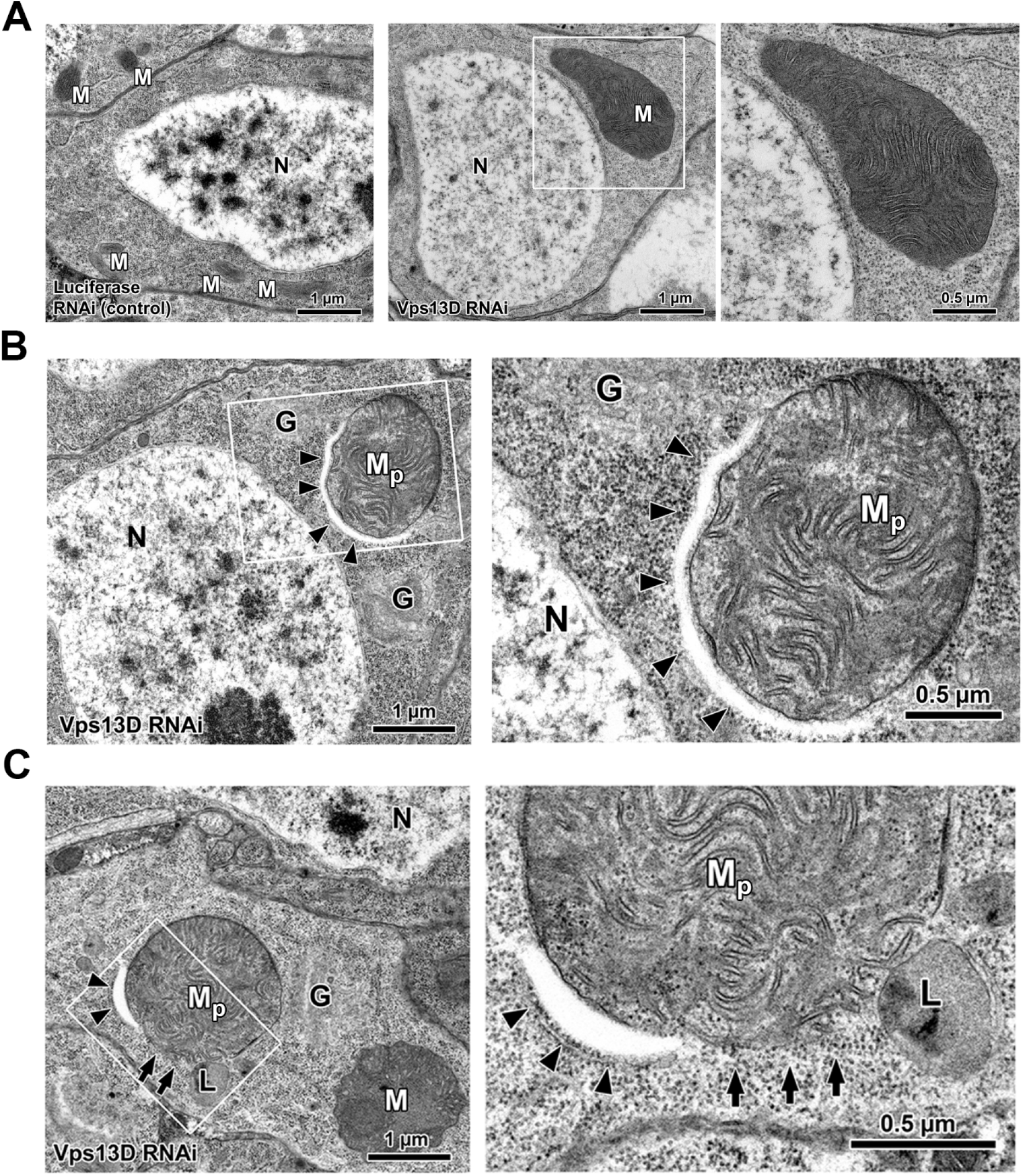
Ultrastructural examination of mitophagy intermediates in neurons depleted for Vps13D. **A)** Representative electron micrographs of larval motoneurons which express RNAi targeting either luciferase (control) or Vps13D, using the pan-neuronal driver Elav-Gal4. Severely enlarged mitochondria (M) in *Vps13D* RNAi condition (right) are readily apparent, in comparison to Control RNAi condition (left). Magnified view of enlarged mitochondria (white box) is shown on the right. Other abbreviations: Golgi (G) and Nucleus (N). **B)** Representative ultrastructural image of a phagophore-associated mitochondria (M_p_) in a *Vps13D*-RNAi depleted neuron. Image on right is a magnified view of cellular region containing the mitophagy intermediate (white box). Arrowheads indicate phagophore. Other abbreviations: Golgi (G) and Nucleus (N). **C)** Representative electron micrograph of a ruptured phagophore-associated mitochondria in a *Vps13D*-RNAi expressing larval neuron. Magnified view of region indicated by the white box is shown to the right. Arrowheads indicate the phagophore; arrows indicate locations lacking mitochondrial membrane, where mitochondrial content appears to mix with the surrounding cytoplasm. Other abbreviations: Mitochondria (M), phagophore-associated mitochondria (M_p_) Golgi (G), and Lysosome (L). Scale bar: 1 μm (left) and 0.5 μm (right).

We never observed mitochondria fully enveloped in what would resemble a completed autophagosome, or localized in lysosomes in micrographs. The incomplete nature of phagophores on mitochondria observed in EM was consistent with three-dimensional rendering of confocal images of Atg8A/B staining suggesting there are indeed many instances of only partially encapsulated mitochondria, while fully encapsulated PolyUb+ mitochondria were never detected in neurons lacking Vps13D (**Figure 2, Supplement 2**). This documented membrane rupture in phagophore-engaged mitochondria in Vps13D depleted neurons provides an attractive explanation for the absence of staining for matrix markers, and suggests that both inner and outer membranes of the mitochondria become disrupted. Because these mitochondrial objects are engaged with a phagophore, we hypothesize that the PolyUb+/Ref(2)p+/ATP5A+/Atg8+/mitoGFP- objects are stalled intermediates in mitophagy. Going forward, we refer to these objects as mitophagy intermediates lacking matrix.

### Mitophagy intermediates lacking matrix are observed in neurons disrupted for Drp1

In *Drosophila* intestinal cells lacking either Vps13D or Drp1, mitophagic clearance was disrupted (Anding et al. 2018), however mitophagy intermediates lacking matrix were not observed. We then wondered whether these neuron-specific intermediates arise in other scenarios of defective mitochondrial fission in neurons. We first utilized Gal4-driven motoneuron expression (D42-Gal4) of *Drp1* RNAi (Sandoval et al. 2014), simultaneous with expression of mitoGFP in motoneurons. Knockdown of Drp1 resulted in extreme enlargement of mitochondria, to a greater degree than loss of Vps13D (**Figure 3, Supplement 1A**), and also led to the presence of Ref(2)p+/ATP5A+/mitoGFP- mitophagy intermediates lacking matrix (**Figure 3A&B)**. Further, these mitophagy intermediates are associated with a phagophore, as shown by the colocalization of ATP5A+/PolyUb+ mitochondria with Atg8A/B (**Figure 3C**) 84.2% of the time.

**Figure 3:**
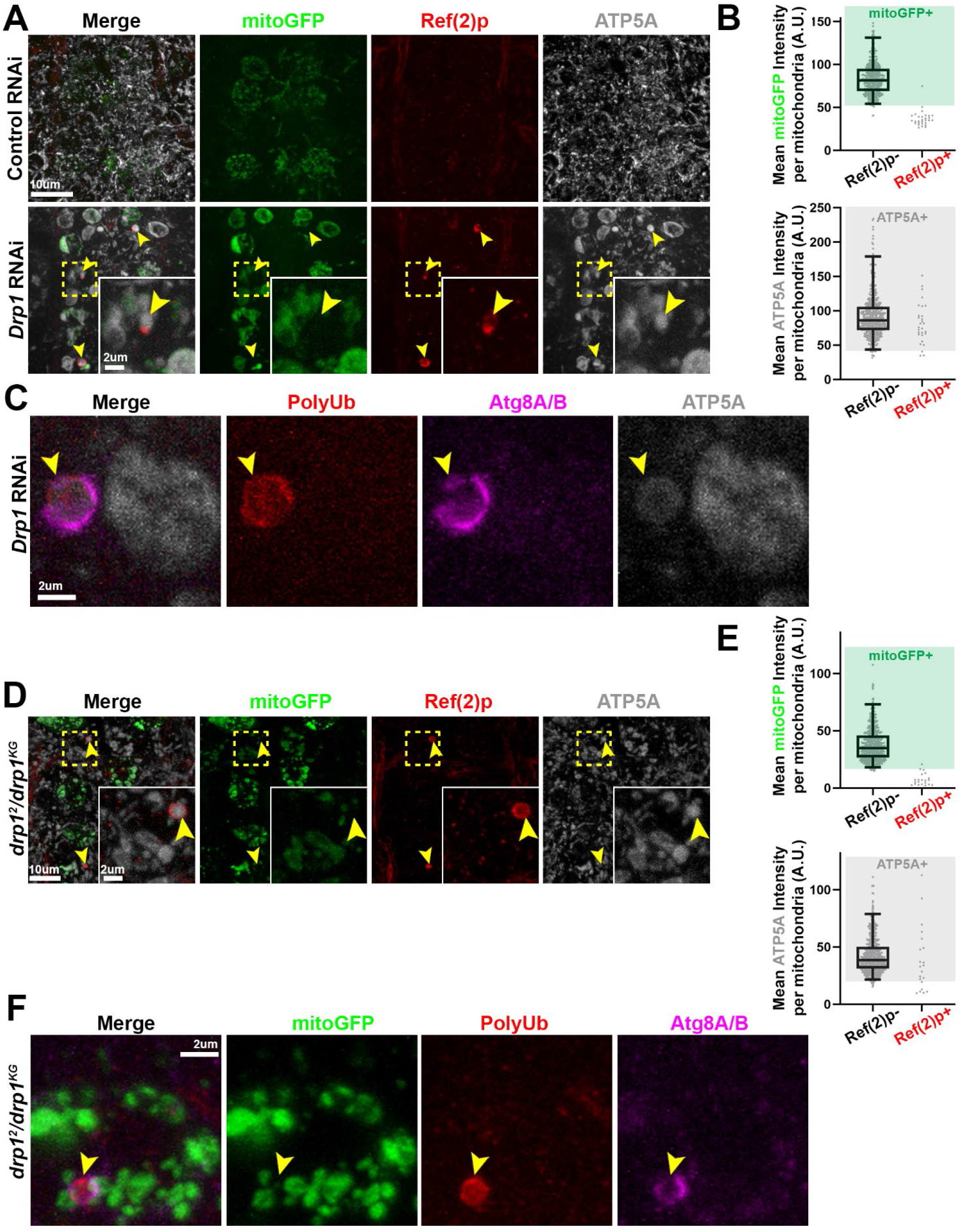
Neurons depleted for Drp1 contain mitophagy intermediates lacking matrix similar to Vps13D depleted neurons. **A)** Representative images of motoneurons in the larval VNC which co-express the mitochondrial marker mitoGFP (green) and RNAi targeting luciferase (control) versus Drp1 (67160), via the D42-Gal4 driver. Tissue was stained for Ref(2)p (red) and ATP5A (greyscale). Dashed yellow box outlines a single Gal4-expressing neuronal cell body that is shown in high magnification in the inset in the bottom right corner. Yellow arrowheads indicate example mitophagy intermediates (Ref(2)p+/ATP5A+/mitoGFP-). **B)** Graph showing the distribution of intensities for mitoGFP (top) and ATP5A (bottom) in conventional mitochondria (ATP5A+/mitoGFP+ and Ref(2)p-), compared to the Ref(2)p+ objects. Based on the criteria of intensity greater than 2.5% of conventional mitochondria (shading), 3.4% of Ref(2)p+ objects contained mitoGFP, while 89.7% contained ATP5A. n=528 mitochondria and 29 PolyUb+ from 8 larval VNCs. See **Figure 1C** and Methods for more detail. **C)** Representative image from a *Drp1*-RNAi expressing neuron which contains two enlarged ATP5A+ mitochondria (greyscale). One, highlighted by the yellow arrowhead, co-localizes with PolyUb (red), and endogenous Atg8A/B (magenta). 84.2% of PolyUb+/ATP5A+ objects co-localize with Atg8A/B (n=38 PolyUb+ mitochondria from 15 larval VNCs). **D)** Representative image of motoneurons in the larval VNC of *drp1^2^/drp1^KG^* mutant larvae, with UAS-mitoGFP expressed via the D42-Gal4 driver. Dashed yellow box outlines a single Gal4-expressing neuronal cell body that is shown in high magnification in the inset in the bottom right corner. Yellow arrowheads indicate example mitophagy intermediates that are postive for Ref(2)p (red) and ATP5A (greyscale), but lack mitoGFP (green). **E)** Graph showing the distribution of intensities for mitoGFP (top) and ATP5A (bottom) in conventional mitochondria (ATP5A+/mitoGFP+ and Ref(2)p-), compared to the Ref(2)p+ objects (n=648 mitochondria and 29 Ref2(p)+ objects from 7 larval VNCs. See **Figure 1C** and Methods for more detail. **F)** Representative image of a mitophagy intermediate (yellow arrowhead) in the VNC of a *drp1^2^/drp1^KG^* mutant larva, with UAS-mitoGFP expressed via the D42-Gal4 driver. The highlighted intermediate in this motoneuron stains for PolyUb (red) and endogenous Atg8A/B (magenta), but lacks the matrix marker mitoGFP. 74.4% of PolyUb+/mitoGFP- objects co-localized with Atg8A/B (n=43 mitophagy intermediates lacking matrix from 8 *drp1* mutant VNCs).

To verify that the mitophagy intermediates are linked to loss of Drp1 function, we examined independent genetic methods to impair Drp1. Based on the mitochondrial enlargement phenotype, the *drp1^KG^* allele (Bellen et al. 2004), which contains a P-element insertion, is a more severe loss of function allele compared to the *drp1^1^* and *drp1^2^* alleles, which contain EMS-generated point mutations (Verstreken et al. 2005). The morphological enlargement of mitochondria in the *drp1^KG^/Df* mutants *(Df = deficiency)* most closely resembled the *Drp1* RNAi conditions **(Figure 3, Supplement 1B)**, but these mutants resulted in lethality during the 2nd instar larval stage. The combination of the morphological and lethality observations suggest that the *drp1^KG^* mutation is a more severe loss of function allele compared to the point mutants, and that expression of the *Drp1* RNAi in neurons results in a phenotype consistent with severe loss of function. Regardless of the allelic combination, we observed the presence of Ref(2)p+/ATP5A+/mitoGFP- mitophagy intermediates in all *drp1* mutants **(Figure 3D&E**, and **Figure 3, Supplement 1C)**, with a frequency that correlated with the severity of the mitochondrial enlargement phenotype. Consistent with other control conditions (Control RNAi and *Vps13D* heterozygous animals), these objects were not observed in *drp1* heterozygous animals (data not shown). Similar to *Vps13D* RNAi, and *Drp1* RNAi conditions, the PolyUb+/mitoGFP- intermediates in *drp1* mutants engage with a phagophore, as they stain positive for Atg8A/B (**Figure 3F**) 74.4% of the time. Overall these results demonstrate that neurons with either disrupted Vps13D or Drp1 contain mitophagy intermediates that uniquely lack mitochondrial matrix, and are engaged with a phagophore.

### Completion of mitophagy differs between the fission-deficient conditions of Vps13D and Drp1 loss

The presence of mitophagy intermediates lacking matrix suggested that mitophagy may be blocked when either Drp1 or Vps13D function is lost. This idea is consistent with previous observations in the cerebellum of *Drp1* knockout mice (Kageyama et al. 2012), However it is has not been tested whether mitophagy is actually blocked; it could alternatively be induced, leading to more detectable intermediates, or slowed, allowing their detection before they are ultimately degraded. We therefore asked whether mitophagy is completed by lysosome-mediated clearance in Drp1 or Vps13D deficient neurons, turning to fluorescent reporters that contain dual acid-labile (GFP) and acid-stable (mCherry) tags to enable selective marking of components that have been successfully trafficked to lysosomes as an estimation of flux. A number of fluorescently tagged mitochondrially targeted reporters have been developed for the purpose of tracking mitophagic delivery to lysosomes (Sun et al. 2015; Cornelissen et al. 2018; Lee et al. 2018), however the loss of matrix proteins from the mitophagy intermediates ruled out the use of matrix targeted reporters such as mitoKiema (Cornelissen et al. 2018). We observed that the OMM targeted mitoQC reporter (Lee et al. 2018; Allen et al. 2013; McWilliams et al. 2016) localized promiscuously in control and Vps13D depleted larval neurons (**Figure 4, Supplement 1A**), in agreement with another recent report (Katayama et al. 2020), and was not consistently present on the Ref(2)p+ mitophagy intermediates in Vps13D depleted neurons (**Figure 4, Supplement 1B**). Hence, the currently available mitochondrially localized reporters in *Drosophila* were not suitable for estimating completion of mitophagy in our genetic conditions.

We therefore turned to a previously characterized mCherry-GFP-Atg8A reporter for autophagy (Klionsky et al. 2016; Takáts et al. 2013; Kimura, Noda, and Yoshimori 2007). We found that this reporter, when stained with GFP antibodies in fixed tissue, co-localized with the Ref(2)p+ mitochondria in Vps13D and Drp1 depleted neurons (**Figure 4, Supplement 2A**), consistent with the endogenous Atg8A/B localization (**Figure 1D**). Dual GFP/RFP-containing phagophores were only detectable in a small subpopulation of Vps13D depleted neurons in live VNC preparations (**Figure 4, Supplement 2B**), likely due to lower fluorescence intensity in active phagophores compared to accumulated signal in lysosomes (Nagy et al. 2015). However, bright red-only signal, indicating the accumulation of reporter delivered to the acidic lysosome compartment, were detectable in the majority of neurons in all conditions in live preparations. The red-only signal was increased in larval motoneurons overexpressing Atg1, which is expected to induce autophagy (Nagy et al. 2015) (**Figure 4, Supplement 3A&B**). Conversely, knockdown of the essential autophagy component Atg5 resulted in less overall red-only signal in neurons containing red-only puncta (38.5% compared to Control RNAi), with a high proportion of neurons lacking red-only signal altogether (47.8%, 34 of 71 neurons) (**Figure 4, Supplement 3A&B**). These results indicate that the total levels of red-only signal in larval motoneurons expressing the mCherry-GFP-Atg8A reporter can represent an estimation of delivery of autophagic substrate to the lysosome.

An experimental condition that has been rigorously demonstrated to induce mitophagy in the *Drosophila* larval nervous system has not been previously established. Based on our observations of mitophagy intermediates in the fission-deficient conditions we tested above, we expected to observe an increased delivery of reporter to lysosomes in both conditions, however we observed strikingly different results between Vps13D and Drp1 depleted neurons. Drp1 depletion led to a strong induction of red-only mCherry-GFP-Atg8A puncta (**Figure 4A&B**, with 0% of neurons lacking red-only puncta). In contrast, Vps13D depleted neurons showed slightly less, though not significantly different, levels of red-only mCherry-GFP-Atg8A signal compared to control neurons (**Figure 4A&B**). We suspect that autophagy levels in Vps13D depleted neurons are indeed lower than control neurons, since in addition to the trending decrease in overall red-only signal, a higher proportion of Vps13D depleted neurons (15.7%, 11 of 70 neurons) lacked red-only signal altogether in comparison to neurons expressing control RNAi (10.4%, 8 of 77 neurons). Regardless of the comparison to control neurons, the difference between Drp1 versus Vps13D depleted neurons is drastic. The observation that either Vps13D or Drp1 depletion result in mitophagy intermediates (**Figures 1 & 3**) but only one, Drp1 depletion, results in increased autophagic delivery to lysosomes suggested that autophagy completion may be blocked in Vps13D depleted neurons.

**Figure 4:**
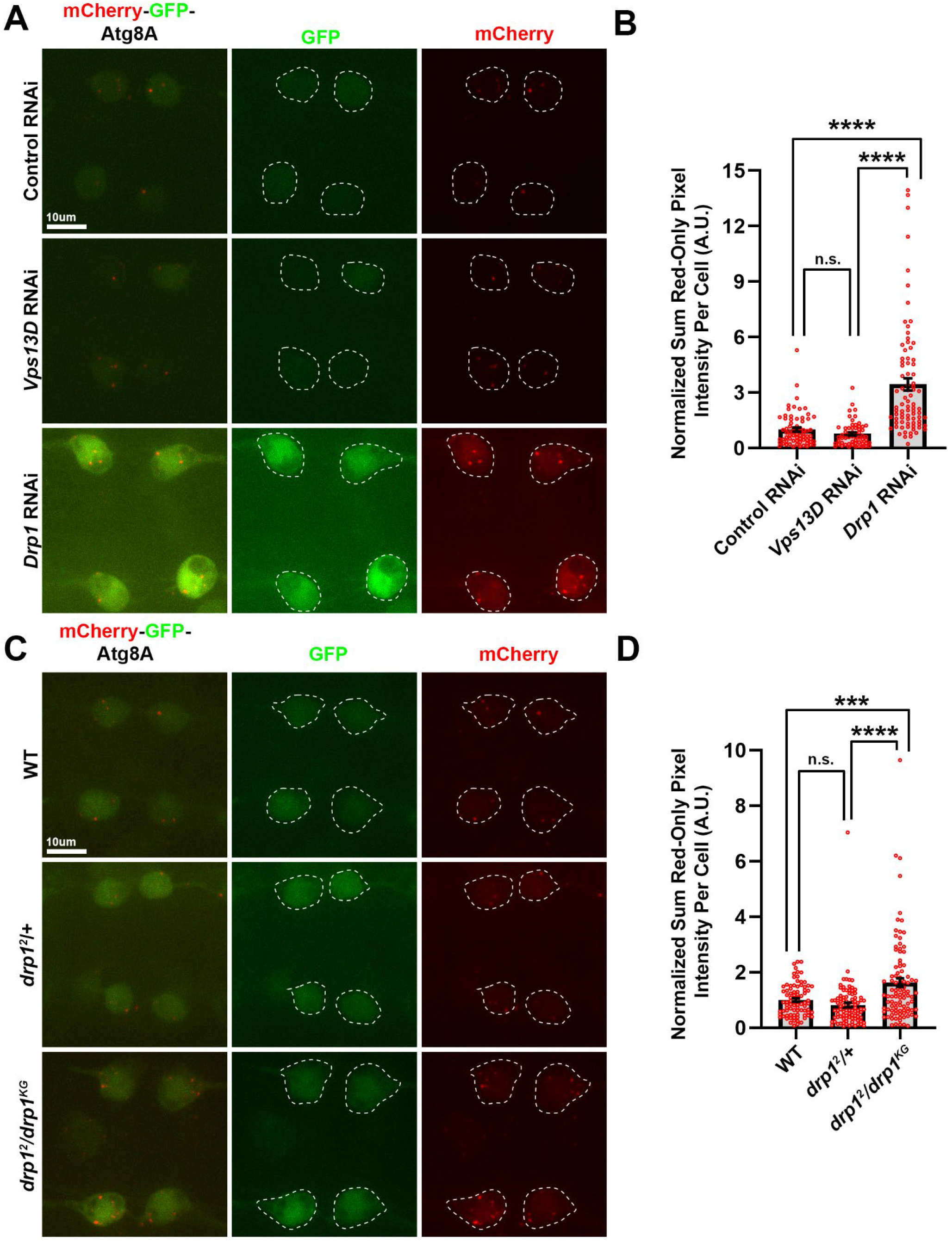
Completion of mitophagy in neurons depleted for Drp1, but not Vps13D. **A)** Representative images of live larval motoneurons expressing UAS-mCherry-GFP-Atg8A, expressed via the D42-Gal4, simultaneous with expression of UAS-RNAi lines targeting luciferase (control) (BL# 31603), *Vps13D* (BL# 38320), or *Drp1* (BL# 67160). White dashed lines indicate the outlines of individual cell bodies. **B)** Quantification of the sum pixel intensity of the red-only signal per neuronal cell body (normalized to Control RNAi). Each point represents a single neuronal cell body, bars represent the mean ± SEM (Control RNAi n= 69 cell bodies; *Vps13D* RNAi n=59 cell bodies; and *Drp1* RNAi n=82 cell bodies from 6 larval VNCs each). **** represents p value <0.0001. n.s. (not significant) represents p value >0.5 (p=0.09). **C)** Representative images of live larval motoneurons expressing UAS-mCherry-GFP-Atg8A, expressed via the D42-Gal4 driver, in the indicated genetic backgrounds. White dashed lines indicate the outlines of individual cell bodies. **D)** Quantification of the sum pixel intensity of the red-only signal per neuronal cell body (normalized to WT). Each point represents a single neuronal cell body, bars represent the mean ± SEM (WT n= 84 cell bodies; *drp1^2^/*+ n= 93 cell bodies; and *drp1^2^/drp1^KG^* n=96 cell bodies from 7 larval VNCs each). *** represents p value <0.001. **** represents p value <0.0001. n.s. (not significant) represents p value >0.5 (p=0.09).

For unknown reasons, despite the same genetic conditions, the expression levels of the mCherry-GFP-Atg8A reporter in the cytoplasm was significantly higher in neurons expressing *Drp1* RNAi compared to any other condition. Since we could not rule out that the increase in red-only puncta is due to the increase in expression levels of the reporter, we examined the mCherry-GFP-Atg8A reporter in motoneurons of *drp1* mutants. Consistent with the knockdown phenotype, autophagic delivery of the tandem-tagged Atg8A reporter to lysosomes is increased in *drp1* mutant neurons as shown by a significant increased in the amount of red-only puncta in mutants compared to WT and heterozygous animals (**Figure 4C&D**). This increase in red-only signal was not associated with increased overall expression of the reporter, as was observed with *Drp1* RNAi expression. The increased red-only signal was suppressed in the condition of Atg5 knockdown (**Figure 4, Supplement 3C&D**), confirming that the increase in signal is dependent upon autophagy machinery. Together these observations suggest that mitophagy intermediates appear in conditions when mitochondrial fission is impaired, however completion of autophagy only occurs in Drp1 depleted or mutant neurons. Completion appears to be stalled in Vps13D depleted neurons. Given the co-localization of the mCherry-GFP-Atg8A reporter with mitochondria in both fission deficient conditions, we interpret this data to indicate that increased mitophagic flux is at least partially responsible for the increased delivery of reporter to lysosomes observed in Drp1 mutant neurons.

### Genetic validation of incomplete mitophagy in Vps13D depleted neurons compared to Drp1 depleted neurons

The interpretation that induction and completion of mitophagy is increased in Drp1 deficient neurons but not Vps13D deficient neurons was originally surprising, so we sought an additional method to assess whether mitophagy is completed in these genotypes. A common technique used in cultured cells is to observe the accumulation of an autophagy substrate when lysosomal degradation is inhibited (Klionsky et al. 2016). For example, in cultured cells the final degradation step can be blocked by inhibiting fusion of the autophagosome with the lysosome using bafilomycin. For an established method to block autophagy in *Drosophila* neurons *in vivo*, we used a null mutation in the core autophagy component *Atg5 (Kim et al. 2016)*, which is required for the initiation and progression of autophagy (Ariosa and Klionsky 2016). If mitophagy is responsible for the differences in autophagic flux observed in Vps13D or Drp1 depleted neurons, then the mitophagy intermediates should build up when Drp1 is knocked down in the background of *Atg5* null mutants. Conversely, if mitophagy is stalled (not completed), then the frequency of mitophagy intermediates should not be affected by the loss of *Atg5*.

We first carried out further analysis of mitochondria and potential mitophagy intermediates in motoneurons of *Atg5* null larvae. While mitochondria in *Atg5* null neurons were slightly enlarged in morphology compared to control animals (**Figure 5, Supplement 1A**), the overall complex network of mitochondria remained intact, and the strongly enlarged and isolated mitochondria (present in Vps13D and Drp1 depleted neurons) were not observed. We observed Ref(2)p+ puncta in *Atg5* null neurons, some of which fully overlapped with ATP5A which were scored as mitophagy intermediates (closed yellow arrowhead, compared to open yellow arrowhead). Ref(2)p+/ATP5A+ species were observed in a relatively low percentage of neurons (7.2%) (**Figure 5A&B**).

**Figure 5:**
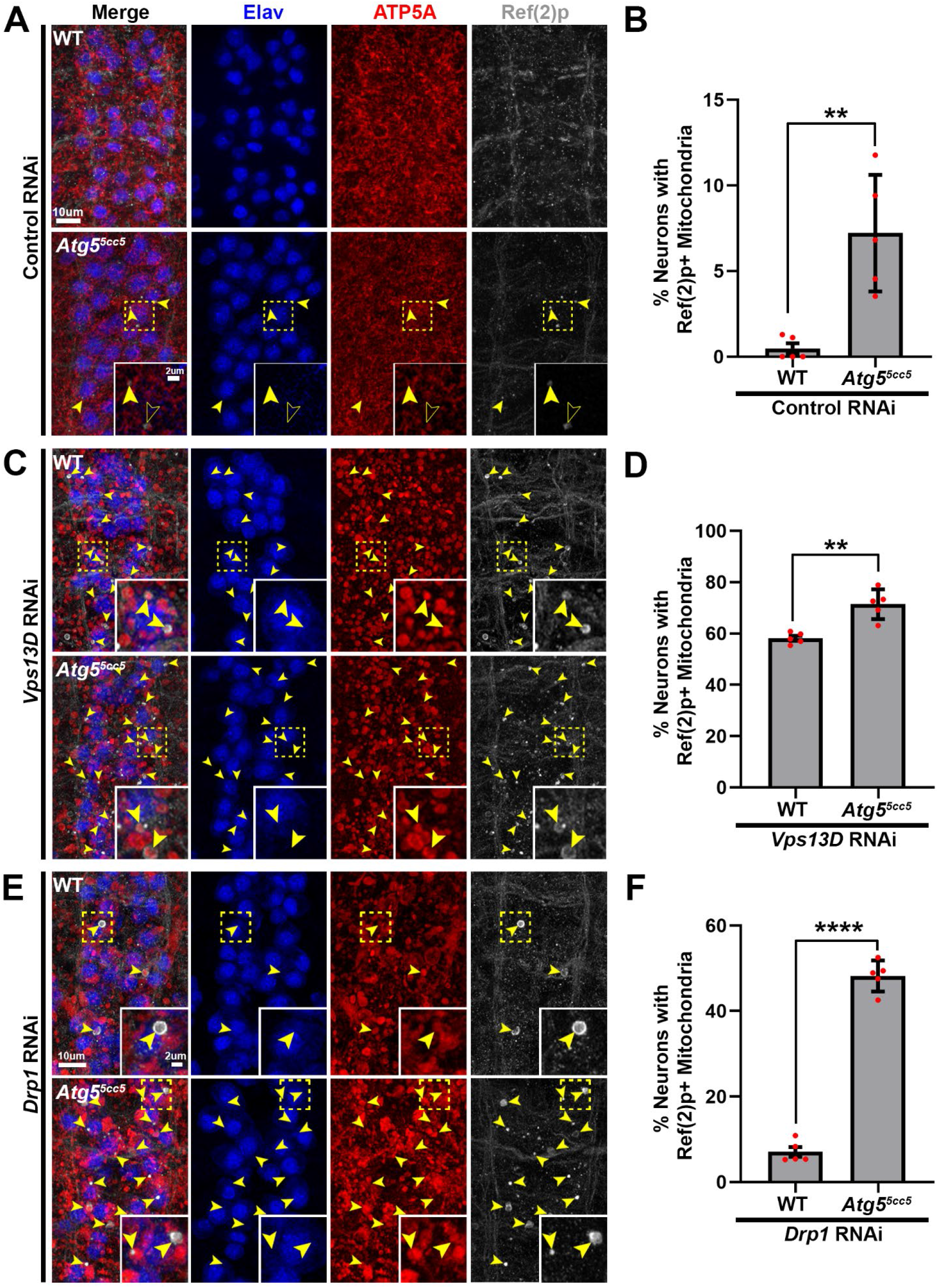
Genetic validation of incomplete mitophagy in Vps13D depleted neurons versus Drp1 depleted neurons. **A,C,E)** Representative images of dorsal midline motoneurons in larval VNCs from a wildtype background (*w^1118^*) (top) or *Atg5* null (*Atg5^5cc5^*) (Kim et al., 2016) background (bottom) expressing (A) UAS-luciferase (Control) RNAi, (C) UAS-*Vps13D* RNAi, or (E) UAS-*Drp1* RNAi. Tissue was stained for neuron-specific transcription factor Elav (blue), mitochondrial protein ATP5A (red), and autophagy receptor protein Ref(2)p (greyscale). Inset in bottom right corner indicates a high magnification image of a single confocal plane from a neuronal cell body (yellow dashed box). Yellow arrowheads indicate mitophagy intermediates (Ref(2)p+/ATP5A+). An additional population of Ref2(p)+ puncta that lacked ATP5A (open arrowheads) were detected in *Atg5* null mutants expressing control RNAi (A). **B,D,F)** Quantification of the percentage of dorsal midline motoneurons containing a mitophagy intermediate (Ref(2)p+ mitochondria). Red points represent the % of neurons from one animal (n=5 animals for each condition). Bars represent mean ± SEM. ** p<0.01; **** p<0.0001.

In *Atg5* null neurons expressing *Vps13D* RNAi, the accumulation of phagophore marker Atg8A/B on PolyUb+ mitochondria was abolished (**Figure 5, Supplement 1C&D**), confirming both the effectiveness of *Atg5* deletion in blocking autophagy and the dependence of Atg8A/B staining on autophagy induction. Importantly, in these conditions, mitochondria were still co-localized with PolyUb, suggesting that they form independently to impairments in autophagy machinery.

We then asked whether blocking autophagy with the *Atg5* mutation led to changes in the number of mitophagy intermediates detected in Vps13D or Drp1 depleted neurons. Loss of *Atg5* led to only minor (1.2 fold increase), but still significant, changes in the number of mitophagy intermediates in Vps13D depleted neurons (**Figure 5C&D**). In contrast, loss of *Atg5* in Drp1 depleted neurons led to a dramatic increase (6.8 fold) in the percentage of neurons that contained Ref2p+ mitophagy intermediates (**Figure 5E&F**). This trend was similarly observed when we knocked down *Atg5* with RNAi in *drp1* mutant neurons (**Figure 5, Supplement 2D**); in this condition we also observed decreased delivery of the autophagy reporter to the lysosome (**Figure 4, Supplement 3C&D**). These observations support the interpretation suggested by the mCherry-GFP-Atg8A reporter data that mitophagy is both induced and completed in Drp1 depleted neurons, however fails to complete in Vps13D depleted neurons (**Figure 6A**).

**Figure 6:**
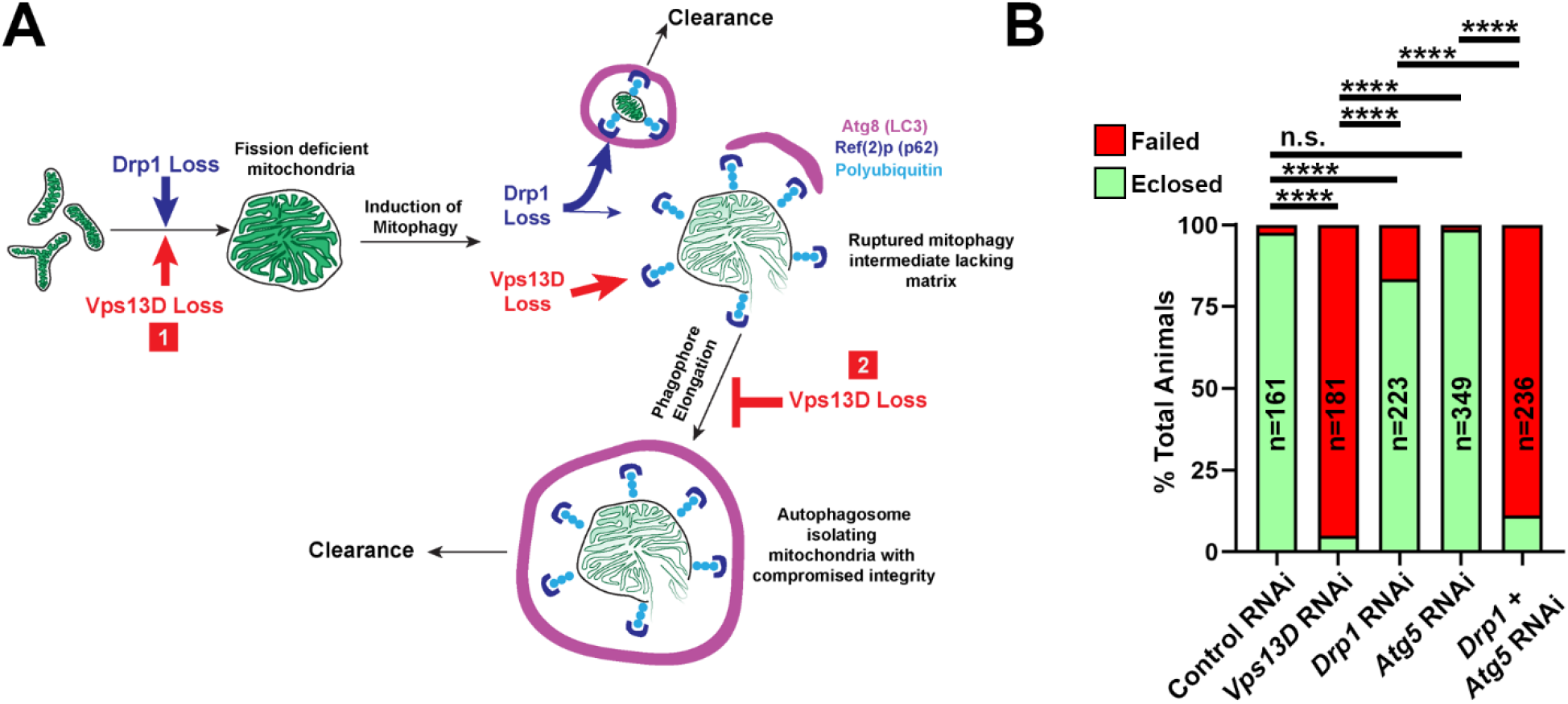
Two-hit model for lethality in Vps13D depleted neurons. **A)** Two-hit model for mitophagy intermediates lacking matrix present in Vps13D depleted neurons knockdown of Vps13D or Drp1. (1) loss of Vps13D and loss of Drp1 lead to similar defects in mitochondrial fission and an induction of mitophagy. In Drp1 depleted neurons (blue arrows and text), mitophagy clears the majority of smaller, matrix-containing mitochondria, while a subpopulation of larger mitophagy intermediates lacking matrix exist that appear stalled. In Vps13D depleted neurons (red arrows and text), a second role for Vps13D is revealed for (2) elongating the phagophore around damaged mitochondria. Therefore, depletion of Vps13D causes the accumulation of mitophagy intermediates (top right) that are polyubiquitinated, coated with Ref(2)p (*Drosophila* homolog of p62), engaged with a stalled phagophore (labeled by Atg8 which is the *Drosophila* homolog of LC3), and lacking in matrix content because of rupture. **B)** Quantification of successful eclosion frequency of pupae with the indicated RNAi driven by pan-neuronal driver nSyb-Gal4. n.s. indicates no significant difference, *** indicates p<0.001, and **** indicates p<0.0001 based on Chi square test of two compared genotypes indicated.

The Ref(2)p+ mitophagy intermediates that become trapped in *Atg5* null; Drp1 depleted neurons, and similarly in *drp1* mutant animals expressing *Atg5* RNAi, showed some significant differences from the intermediates lacking matrix that have been described thus far: they are smaller in size, largely contain mitoGFP (**Figure 5, Supplement 2**). Therefore, the lack of mitochondrial integrity and loss of matrix in mitophagy intermediates is not a default consequence of impaired mitophagy completion. We interpret that Drp1 loss leads to the induction of two distinct classes of mitophagy intermediates. The first retain their matrix and are usually undetectable due to rapid destruction by autophagy. The second, less abundant, class of intermediates lose their matrix and, similarly to the intermediates that accumulate in Vps13D mutants, may be stalled for mitophagy completion since their numbers do not increase upon loss of Atg5.

### Functional importance of mitochondrial dynamics and mitophagy in neurons

We expected that the failed clearance of mitophagy intermediates in Vps13D and Atg5; Drp1 depleted neurons should have negative consequences for neurons and the overall health of animals. While we observed no evidence for apoptosis, we noticed that animal survival negatively correlates with the accumulation of mitophagy intermediates. Pan-neuronal (via nSyb-Gal4) knockdown of Vps13D leads to severe lethality in flies; only 5% of animals successfully eclosed from their pupal cases. In comparison, knockdown of Drp1 using the same Gal4 driver leads to a successful eclosion rate of 84% (**Figure 6B**). Simultaneous neuron-specific knockdown of the core autophagy protein Atg5 along with Drp1 causes a significant decrease in eclosion rate (11%), which is much lower than in the cases of knocking down either protein alone (**Figure 6B**). These observations are consistent with a two-hit model (**Figure 6A**) for toxicity: loss of Drp1 alone is only mildly toxic because induction of mitophagy can clear defective mitochondria. In addition, loss of autophagy alone is not toxic because neurons undergo only low levels of basal mitophagy in the absence of any additional stressors. However, scenarios that both induce mitochondrial damage and impair clearance lead to strong lethality. Impairments in Vps13D function can lead to dual defects in damage and defective clearance, providing an attractive explanation for the accelerated neurological pathophysiology associated with patients harboring mutations in this gene (Seong et al. 2018; Gauthier et al. 2018; Koh et al. 2020).

## Discussion

Since the discovery of a role for PD associated proteins Parkin and PINK1 in removal of damaged mitochondria via mitophagy, there has been a sustained effort to understand the importance of mitophagy in the maintenance of neuronal health and function. However, discoveries made from mitophagy assays performed in cultured cells have been difficult to verify *in vivo* due to the lack of a robust means of detecting, experimentally inducing, or blocking mitophagy in neurons. Detection of mitophagy, even in cultured cells, generally requires an increase in mitochondrial damage from a stress to drive mitochondria into mitophagy, or the blockage of mitophagy to visualize mitochondria stalled in the pathway. Our data suggest that Vps13D depletion in neurons does both of these, which leads to the accumulation of stalled mitophagy intermediates (**Figure 6A**). First, loss of Vps13D disrupts fission; this stress leads to the induction of mitophagy. Second, loss of Vps13D disrupts mito-phagophore elongation; this leads to the accumulation of stalled mitophagy intermediates. Lastly, stalled mitophagy intermediates in Vps13D depleted neurons lose integrity, rupture, and lose their matrix content. The identification of these intermediates and their readily visualizable features (including their large size) opens up future opportunities to study the molecular mediators and mechanism of neuronal mitophagy.

While the genetic manipulations we performed here may represent extreme experimental conditions, the two-hit defect of accumulated mitochondrial damage combined with diminished capacity for clearance is considered to occur naturally during aging and to be accentuated by conditions of stress and/or genetic mutations in many neurological diseases (Pickrell and Youle 2015). Our ability to dissect these causally related defects in the study of Vps13D should promote new inroads into understanding the pathophysiology of neurological disease.

### Mitochondrial fission in neurons

At face value, the finding that mitophagy can occur when Drp1 function is impaired may be surprising, since several studies have reported an essential role for mitochondrial fission in mitophagy (Twig et al. 2008; Tanaka et al. 2010; Frank et al. 2012). In addition, Drp1 and Vps13D mutants were reported to show equivalent defects in mitochondrial clearance in *Drosophila* intestinal cells (Anding et al. 2018). However, other studies have documented fission-independent mitophagy (Yamashita et al. 2016). We propose that differences in these observations may relate to differences between mitophagic clearance and quality-control mitophagy, and differences in cell types. Fission may be important for breaking down a highly fused tubular mitochondrial network during mitophagic clearance (Rambold et al. 2011). However, neurons are highly reliant on their mitochondria and are not expected to remove all of their mitochondria; instead, mitophagy of only the most damaged mitochondria for quality-control is expected to be critical for long term maintenance of a healthy pool of mitochondria. Our findings imply that neurons deficient for Drp1 and/or Vps13D function lead to mitochondrial damage that induces mitophagy. While stalled mitophagy intermediates have been observed in Drp1-ablated Purkinje cells in the mouse cerebellum (Kageyama et al. 2012), the interpretation of the existence of these intermediates differed for ours, since it was inferred that mitophagy was blocked. In contrast, our findings indicate that mitophagy is induced and completed in Drp1 mutant *Drosophila* neurons, while a small and unique population of mitochondria lacking matrix (which may be ruptured), appear stalled as mitophagy intermediates.

How could disruption of mitochondrial fission induce mitophagy in neurons? Previous studies have shown that conditional genetic ablation of Drp1 in either mouse cerebellar or hippocampal neurons leads to oxidative stress with varying outcomes (Kageyama et al. 2012; Oettinghaus et al. 2016) (cell lethality or the induction of Nrf2-dependent stress response, respectively). Accumulation of mitochondrial reactive-oxygen species (ROS) has previously been shown to be a trigger of mitophagy (Xiao et al. 2017). It will therefore be interesting future work to investigate the potential contributions of mitochondrial ROS in Vps13D mutant neurons in animals and ataxia patients, as administration of antioxidant compounds was capable of suppressing the neurodegenerative phenotype associated with Drp1 loss in Purkinje cells (Kageyama et al. 2012).

We propose that accelerated mitochondrial damage caused by disruptions in fission lead to increased mitophagy in neurons as an adaptive response required to efficiently degrade damaged organelles. Our results suggest that mitophagy intermediates pose a cellular threat, as shown by the severe lethality caused by either Vps13D loss in neurons and the combined depletion of Drp1 and Atg5 (**Figure 6B**). Other reports have documented increased induction of autophagy upon disruptions of the balance of mitochondrial fission and fusion in neurons including *Drosophila* models of Charcot-Marie Tooth Type 2A syndrome (CMT2A) (El Fissi et al. 2018), and rodent models of autosomal dominant optic atrophy (Zaninello et al. 2020).

### Mitochondrial rupture in Vps13D depleted neurons

The rupturing of mitochondrial membranes in Vps13D depleted neurons observed in EM images of stalled mitophagy intermediates was consistent with our staining showing a lack of mitochondrial matrix in mitophagy intermediates. Our observation that mitoGFP does not label stalled mitophagy intermediates in Vps13D or Drp1 depleted neurons (**Figure 1** and **Figure 3**) serves as a cautionary note that widely-used matrix-targeted fluorescent reporters (Sun et al. 2015; Cornelissen et al. 2018) may fail to detect some forms of mitophagy intermediates, hence may lead to inaccurate interpretations about the state of mitophagy in some scenarios.

Other reports have demonstrated rupture of the outer mitochondrial membrane (OMM) during toxin-induced bulk mitophagy in cultured cells, which enables the phagophore to interact with autophagy receptors, such as prohibitin 2, in the IMM (Yoshii et al. 2011; Chan et al. 2011; Wei et al. 2017). In contrast, our observations document rupture of both the OMM and IMM membranes. Mitochondrial swelling and rupture has been observed in many cell types undergoing various forms of cell death, which is typically mediated by a so-called mitochondrial permeability transition pore (MPTP) (Kinnally et al. 2011). However, our ultrastructural analysis and cleaved caspase staining did not reveal any dying cells in the VNC at the larval stages when we deplete Vps13D from neurons (data not shown). We also note that the ruptured mitochondria lacking matrix do not appear vacuolated, as has been described in other neurodegenerative disease models (Bendotti et al. 2001; Higgins, Jung, and Xu 2003; Kong and Xu 1998).

To our knowledge, this is the first direct documentation of such clear rupturing of mitochondria undergoing mitophagy in neurons *in vivo*. We hypothesize that the ruptured mitochondria are responsible for the strong pupal lethality of neuronal Vps13D depletion. While we did not observe direct evidence of neuronal apoptosis, we consider that mitochondrial DNA and oxidized mitochondrial proteins are potent inducers of innate immune responses (West 2017), and neuro-immune interactions are increasingly being recognized as important factors in neurodegenerative disease (Picca et al. 2020; Yu et al. 2020). Therefore, there is interesting work ahead to understand the pathology that evolves following Vps13D loss.

### Vps13 proteins and cellular function

The stalling of phagophores that we have observed in Vps13D depleted neurons appears most similar to phenotypes that have been described for *Atg2* mutants in yeast and flies (Lu et al. 2011; Velikkakath et al. 2012; Nagy et al. 2015). Importantly, Atg2 contains a unique Chorein_N domain that exists only in Atg2 and in members of the Vps13 protein family. Recent work has demonstrated that this domain, from the Vps13 family member in *C. thermophilum*, can directly function as a phospholipid transporter *in vitro* (Kumar et al. 2018; P. Li et al. 2020; Prinz and Hurley 2020). Taken together, these observations raise an interesting hypothesis that Vps13D, and potentially other Vps13 family members, function similarly to Atg2 to enable phospholipid transport critical for phagophore elongation in neurons. This idea is supported by previous findings that the *Dictyostelium* homolog of Vps13A/C is required for proper autophagic flux (Muñoz-Braceras, Calvo, and Escalante 2015). Important future questions include: do all Vps13 family members have a similar function phagophore elongation? And are these functions separable from each other and that of Atg2? One idea is that these proteins may be used for similar functions but in different contexts, for instance mitophagy versus bulk autophagy. Interestingly, loss of Vps13D in larval fat bodies does not disrupt starvation-induced autophagy, suggesting a specialized function in some but not all forms of autophagy (Anding et al. 2018).

Vps13D’s role in mitochondrial fission may also rely on its function in inter-organelle phospholipid transport or communication, since contacts between the endoplasmic reticulum (ER) and mitochondria have been shown to play a direct role in fission (Friedman et al. 2011), as proper phospholipid composition in mitochondrial membranes modulates Drp1 function (Adachi et al. 2016).

Do the other Vps13 family members share Vps13D’s dual roles in mitochondrial fission and phagophore elongation? Morphological defects in mitochondria have been noted in cell lines lacking Vps13A (Yeshaw et al. 2019), and cells lacking Vps13C (Lesage et al. 2016). However, loss of Vps13D is uniquely lethal in animal models (Seong et al. 2018; Anding et al. 2018; Vonk et al. 2017). We hypothesize this may reflect an essential role for Vps13D in mitochondrial fission in all cell types. In neurons, fission defects lead to an induction of mitophagy, which reveals Vps13D’s second function in phagophore elongation. With this new method to induce mitophagy/autophagy in neurons, future work can evaluate whether other Vps13 family members are also required for mito-phagophore elongation.

In conclusion, we have shown that Vps13D is essential for two processes that are critical to general mitochondrial health: mitochondrial fission and mitophagy (**Figure 6A**). This knowledge now establishes a framework for future work to determine whether patient mutations in *VPS13D* differentially affect one or both of these processes, which should lead to a better understanding of disease pathogenesis in patients with mutations in this gene (Seong et al. 2018; Gauthier et al. 2018; Koh et al. 2020).

## Materials and Methods

### Fly maintenance and Drosophila stocks

Fly stocks were maintained on standard Semi-defined yeast-glucose media. All experiments used feeding (not wandering) 3rd instar larvae which were cultured at 25° in a 12:12h light:dark cycle incubator.

The following strains were used in this study (“BL” indicates a strain from Bloomington Stock Center): *Vps13D* RNAi (BL #38320), *luciferase* RNAi (Control RNAi) (BL #31603), D42-Gal4 (BL #8816), Elav-Gal4 (BL #458), *Vps13D* mutants (*Vps13D^11101^* from BL #56282, and *Df* for *Vps13D* BL #25688), *Atg5* mutant (*Atg5^5cc5^* from G. Juhasz’s lab (Kim et al. 2016)), *Drp1* RNAi (BL #67160), *Atg5* RNAi (BL #34899), *drp1* mutants (*drp1^1^* from BL #24885; *drp1^2^* from BL #24899; *drp1^KG^* from BL #13510; *Df* for *drp1* BL #7494), UAS-mCherry-GFP-Atg8A (gift from T. Neufeld lab), UAS-mitoQC (received from A. Whitworth lab, (Lee et al. 2018)), UAS-Idh3a-HA (gift from I. Duncan lab (Duncan, Kiefel, and Duncan 2017)), and nSyb-Gal4 (BL #51635).

An unbiased mixture of both male and female larvae were selected for all experiments unless listed below. Because of the presence of the *Atg5* gene on X-chromosome, experiments in **Figure 5** were exclusively performed in male larvae to achieve homozygosity (*Atg5^5cc5^*/y).

### Immunohistochemistry

Third instar larvae were selected based on visual and fluorescent markers, and dissected in ice-cold PBS. Fixation was either done with 4% formaldehyde in PBS for 20 minutes at room temperature, or with undiluted Bouin’s Fixative (Ricca Chemical Cat# 1120-16) for 7 minutes at room temperature. Following fixation, dissected larvae were washed extensively in PBS, followed by PBS-T (PBS with 0.1% Triton X-100) before blocking for at least 30 minutes at room temperature in PBS-T supplemented with 5% normal goat serum. Primary and secondary antibodies were diluted in the same buffer used for blocking. Primary antibodies were incubated overnight at 4°, followed by PBS-T washes. Secondary antibodies were incubated for at least 2 hours at room temperature, followed by PBS-T washes. Following staining, filleted larvae were mounted using Prolong Diamond mounting media (Life Technologies).

The following primary antibodies were used in this study: anti ATP5A* at 1:1000 (Abcam #ab14748), anti dsRed at 1:500 (Takara #632496), anti Polyubiquitin (“PolyUb”) at 1:50 (FK1) (Enzo Life Sciences #pw8805), anti Ref(2)p at 1:500 (Abcam #ab178440), anti Atg8 (GABARAP)* at 1:500 (Cell Signaling Technology #13733S), anti GFP at 1:1000 (Aves Labs #GFP1020), anti GFP at 1:1000 (Life Technologies #A6455), anti GFP** at 1:500 (Life Technologies #A-11120), anti PDH (PDH-e1ɑ)** at 1:200 (Abcam #ab110334), anti Multi Ubiquitin (“Ub/PolyUb”) at 1:200 (FK2) (MBL #D058-3), anti Elav at 1:100 (Developmental Studies Hybridoma Bank #Rat-Elav-7E8A10), and anti HA at 1:500 (Sigma #H6908). Antibodies designated with the (*) symbol were only used in conditions in which tissue was fixed with Bouin’s Fixative due to dramatically better staining in this fixation condition, whereas antibodies labeled with the (**) symbol were only used in conditions in which tissue was stained with 4% formaldehyde due to dramatically better staining in this fixation condition. Any antibodies not designated with those symbols worked well in both fixation conditions.

The following secondary antibodies (all derived from goat, diluted 1:1000) were used (all from Life Technologies): anti Rabbit (Alexa Fluor 405/488/568/647), anti Mouse IgG1 (Alexa Fluor 568/647), anti Mouse IgG2a (Alexa Fluor 488), anti Mouse IgG2b (Alexa Fluor 488/647), anti IgM (Alexa Fluor 568), and anti Rat (Alexa Fluor 488/647).

### Electron Microscopy

Third instar larvae were dissected in ice cold PBS then fixed with 3.2% paraformaldehyde, 1% glutaraldehyde, 1% sucrose and 0.028% CaCl_2_ in 0.1 N sodium cacodylate (pH 7.4, overnight, 4 °C). Samples were then postfixed with 0.5 % OsO_4_ (1h, RT) then half-saturated aqueous uranyl acetate (30 min, RT), dehydrated in graded series of ethanol and embedded using Spurr Low-Viscosity Embedding Kit (EM0300, Sigma-Aldrich) according to the manufacturer's instructions. Ultrathin 70-nm sections were stained in Reynold’s lead citrate and viewed at 80kV operating voltage on a JEM-1011 transmission electron microscope (JEOL) equipped with a Morada digital camera (Olympus) using iTEM software (Olympus).

### Imaging and Quantification

Confocal images were collected on an Improvision spinning disk confocal system, consisting of a Yokagawa Nipkow CSU10 scanner, and a Hamamatsu C1900-50 EMCCD camera, mounted on a Zeiss Axio Observer with 63X (1.4NA) oil objectives or a Leica SP5 Laser Scanning Confocal Microscope with a 63x (1.4NA) oil objective. Images in **Figure 2, Supplement 2** of individual mitophagy intermediates were subjected to deconvolution using Leica LAS AF deconvolution module. Similar settings were used for imaging of all compared genotypes and conditions. Volocity software (Perkin Elmer) was used for intensity measurements and quantification of all confocal data. All images from conditions of pan-neuronal manipulations (D42-Gal4 or Elav-Gal4) were taken and quantified from the dorsal midline motoneurons (approximately segments A3-A7).

For live larval imaging, third instar larvae were dissected in room temperature HL3 (Stewart et al. 1994) with 0.45mM Ca^2+^, as previously described (J. Li et al. 2017). Snapshots of neurons from larval VNCs were captured with a 63x objective lens using the Improvision spinning disk confocal system. All images acquired in conditions that were normalized together were obtained with identical microscope settings.

To quantify the mean intensities of mitochondrial proteins in PolyUb+ or Ref(2)p+ objects (**Figure 1C; Figure 1, Supplement 2B; Figure 3B&D**) (mitoGFP, ATP5A and PDH), the average intensity of mitochondrial protein staining was collected from >500 individual *bona fide* mitochondria that were PolyUb-. Objects with intensities above the 2.5% percentile of this population (indicated by the semi-transparent rectangles) were scored as positive.

To quantify the phagophore engulfment of mitochondria (**Figure 2, Supplement 2**), separate segmented populations were defined based on intensity using the Volocity software for PolyUb and Atg8 for each mitophagy intermediate, along with a third population which represented the voxel space occupied by the overlap of these two populations. The overall accumulation of Atg8 staining (volume (um^3^) multiplied by the intensity (A.U.)) was plotted against the voxel overlap of the two populations (Atg8 and PolyUb) to test for a correlation. The expectation is that there would be a positive correlation between the size of the phagophore and the overlap of Atg8 and PolyUb signal if the phagophore was successfully engulfing the PolyUb+ mitochondria.

To quantify the red-only signal from live imaging of neurons expressing mCherry-GFP-Atg8A (**Figure 4** and **Figure 4, Supplement 3**), neurons with reporter expression were identified by their diffuse, cytoplasmic localization of the reporter (typically in the GFP channel). For each neuron, a selection algorithm was designed in Volocity to select bright, red-only signal, and similarly applied to all individual neurons. The total red-only signal, accounting for the brightness and total amount (sum pixel intensity), was normalized to 1 for control conditions that were carried out in parallel for each experiment. Conditions that were compared were subjected to the same imaging settings. If a neuron did not contain any red-only signal, it was excluded from comparison to other conditions, but the % of neurons lacking red-only signal was noted for each condition.

To quantify the % of polyubiquitinated mitochondria with phagophores (in text and **Figure 5, Supplement 1C&D**), ATP5A+/PolyUb+ objects were individually assessed for the presence of recognizable Atg8 staining.

To quantify the % of neurons containing mitophagy intermediates (**Figure 5**), mitophagy intermediates were identified as Ref(2)p+ objects that contained ATP5A, and were confirmed to be neuronal based on proximity to labeling of the neuronal nuclear marker Elav. The number of neurons containing one or more mitophagy intermediates were counted from the entirety of motoneurons in the dorsal midline from each ventral nerve cord.

Volume measurements of mitophagy intermediates (**Figure 6, Supplement 1A**) were performed in Volocity, and based off of the volume of Ref(2)p+, ATP5A+ objects.

### Eclosion Assay

Pupae of the proper genotype (based on morphology and fluorescence) were counted 8 days following egg laying, and counted pupae were tracked for a total of 7 days. Successful eclosion was counted as complete exit from the pupal case, as flies that died while partially eclosed were counted as failed.

### Statistics

All statistical methods utilized are listed in the Figure legends, and in almost all scenarios (except for correlation test in **Figure 2, Supplement 2** and eclosion assays in **Figure 6B**) Two-tailed unpaired T-tests assuming parametric distributions were used to compare different conditions. All error bars represent standard error of the mean (SEM), and individual data points are included to indicate the distribution of the data. The statistical significance of Eclosion Assays were determined using a Chi-Square Test to compare two individual genotypes at a time. Sample sizes were determined based on previous literature.

## Acknowledgments

We would like to thank all members of the Collins lab for helpful discussions on this manuscript, and Dr. Margit Burmeister for initiating this collaboration on Vps13D. We thank Eric Robertson for technical assistance with *Drosophila* stock maintenance, and Monika Truszka for technical assistance during EM sample preparation. This work is funded by the following grants: R21NS107781-01A1 and R01NS069844-09 to C.A.C.; K99NS111000-01 to R.I.; KKP129797 and GINOP-2.3.2-15-2016-00032 to G.J.; and PPD-222/2018 (Hungarian Academy of Sciences) to P.L.

## Author Contributions

This work was conceived by R.I. and C.A.C. With the exception of the circumstances listed below, all experiments, data collection, analysis, and interpretation was carried out by R.I. All electron microscopy experiments (Figure 2) were carried out by P.L. and G.J. Useful reagents were also contributed from the lab of G.J. A.W. contributed to eclosion experiments in Figure 6. The manuscript was written by R.I. and C.A.C., and reviewed by all authors.

---

**Figure 1, Figure Supplement 1:**
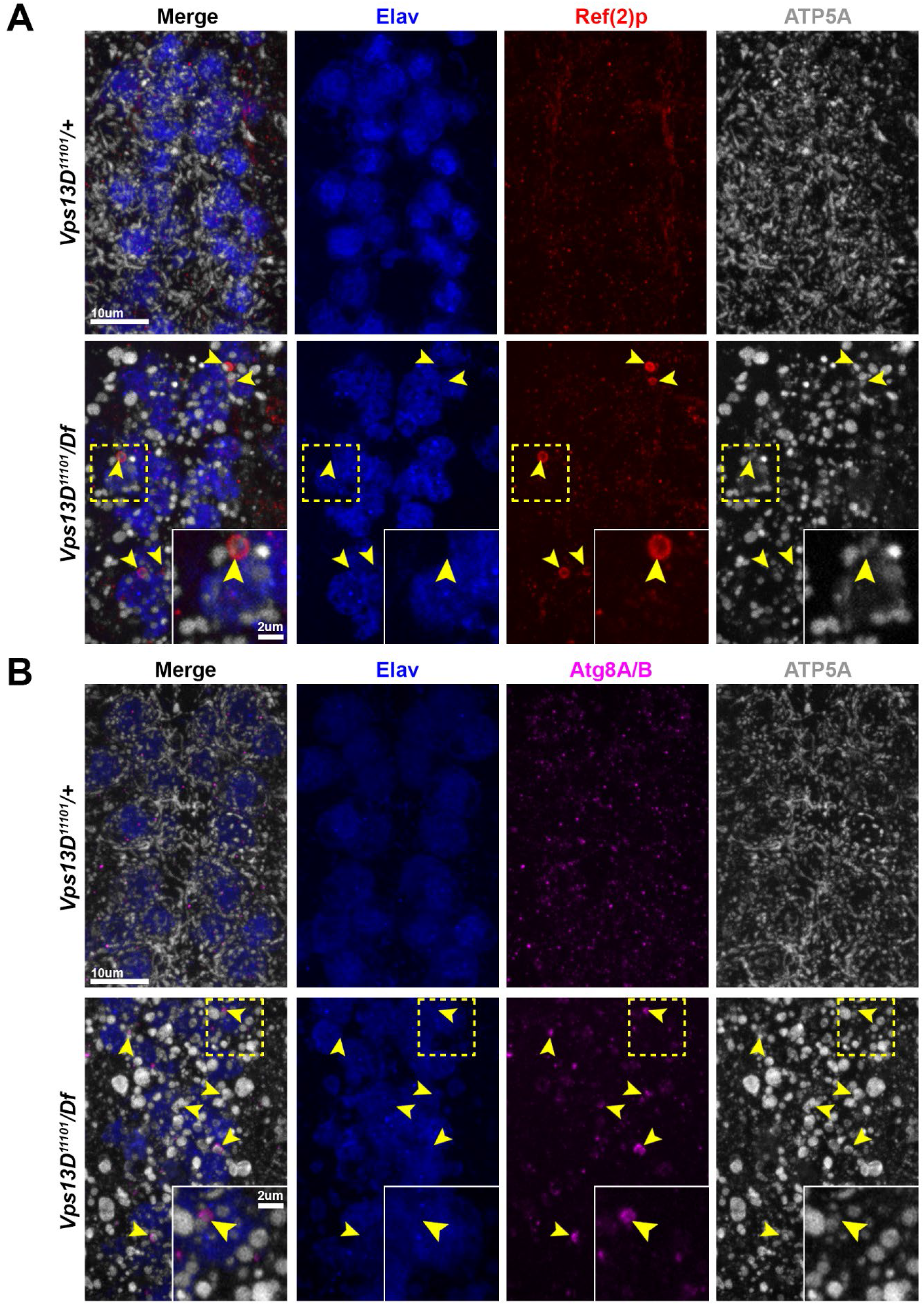
*Vps13D* mutant larvae contain mitophagy intermediates in neurons similar to RNAi condition. **A)** Representative images of dorsal midline motoneurons in early second instar larval VNCs from a *Vps13D* heterozygous (*Vps13D^11101^/+)* background (top) or *Vps13D* mutant (*Vps13D^11101^/Df)* background (bottom). Tissue was stained for the neuron-specific transcription factor Elav (blue), autophagy receptor protein Ref(2)p (red), and mitochondrial protein ATP5A (greyscale). Yellow arrowheads indicate mitophagy intermediates (Ref(2)p+/ATP5A+). Inset in bottom right corner indicates a high magnification image of a single neuronal cell body (yellow dashed box) containing a mitophagy intermediate. **B)** Representative images of dorsal midline motoneurons in early second instar larval VNCs from a *Vps13D* heterozygous background (top) or *Vps13D* mutant background (bottom). Tissue was stained for neuron-specific transcription factor Elav (blue), phagophore marker Atg8A/B (magenta), and mitochondrial protein ATP5A (greyscale). Yellow arrowheads indicate mitophagy intermediates engaged with a phagophore (Atg8+/ATP5A+). Inset in bottom right corner indicates a high magnification image of a single neuronal cell body (yellow dashed box) containing a mitophagy intermediate.

**Figure 1, Figure Supplement 2:**
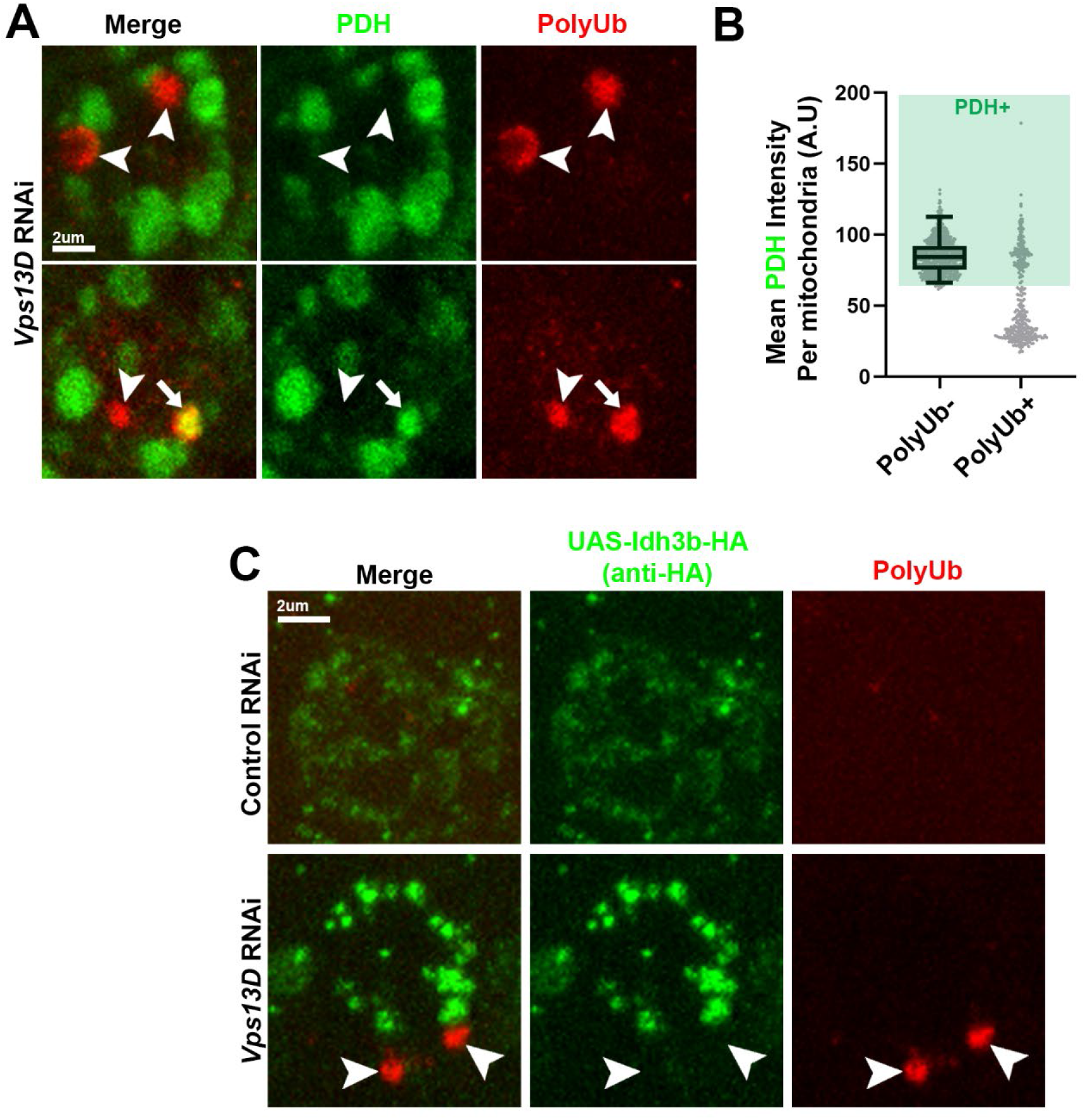
Additional mitochondrial matrix markers are lacking in PolyUb+ objects in Vps13D depleted neurons. **A)** Representative images of single motoneuron cell bodies which express *Vps13D* RNAi via the D42-Gal4 driver, which are stained for endogenous mitochondrial matrix protein pyruvate dehydrogenase E1ɑ (PDH), and PolyUb (red). White arrowheads indicate examples of PolyUb+ objects lacking PDH staining; the white arrow indicates an example of a PolyUb+ object that contains PDH. **B)** Graph showing the distribution of mean PDH intensities conventional mitochondria (PolyUb-) compared to the PolyUb+ objects. Based on the >2.5% criteria (shading), 33.6% of the PolyUb+ objects contain PDH. n=789 mitochondria and 351 polyUb+ objects from 5 larval VNCs. **C)** Representative images of individual dorsal midline motoneuron cell bodies which co-express UAS-luciferase-RNAi (control) versus UAS-*Vps13D* RNAi together with the full-length tagged matrix protein Idh3b-HA (UAS-Idh3b-HA), driven by the pan-motoneuron driver D42-Gal4. Idh3b is detected based on antibody staining for the HA tag (green). White arrowheads indicate PolyUb+ (red) mitophagy intermediates that lack Idh3b-HA.

**Figure 2, Figure Supplement 1:**
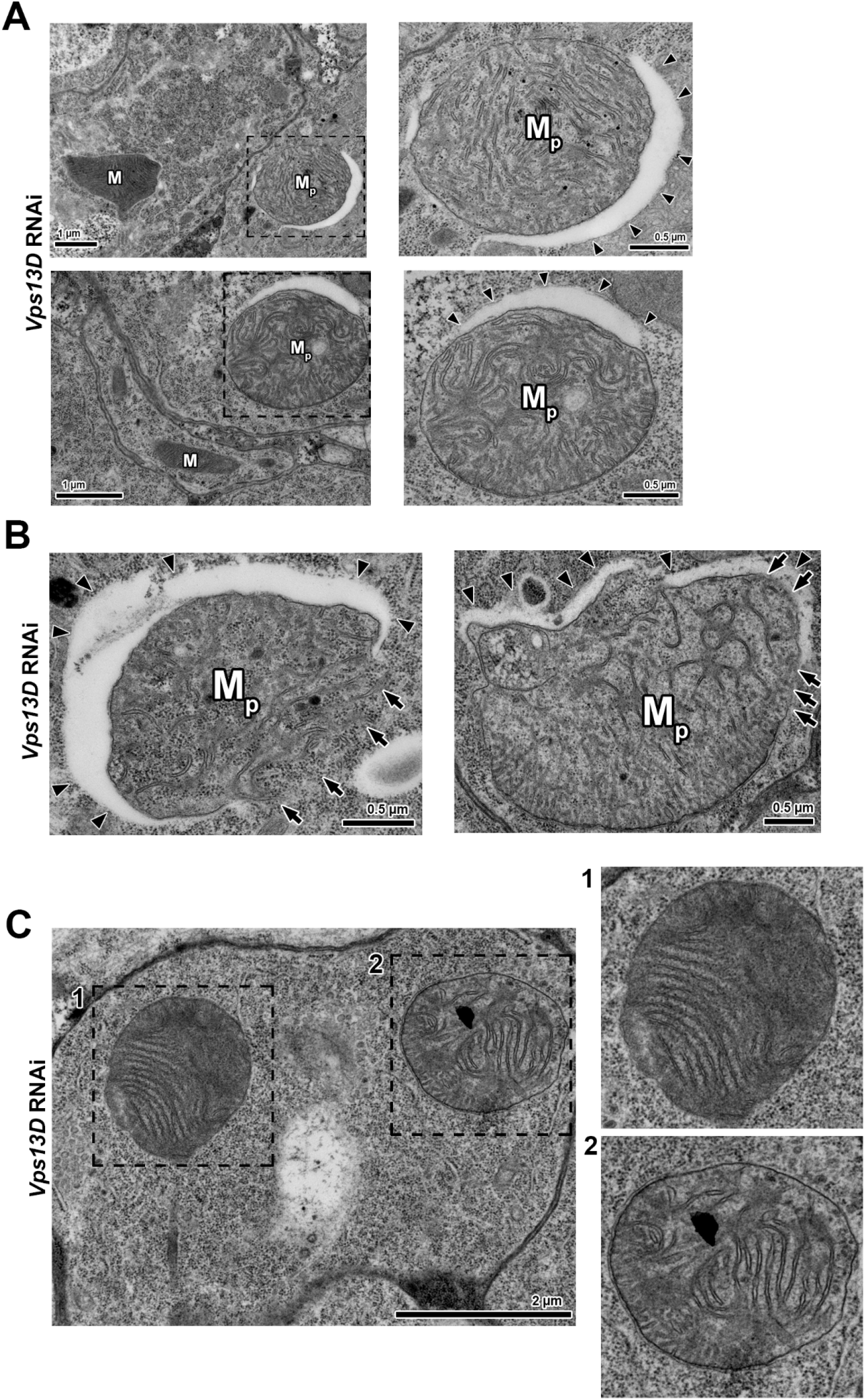
Additional examples of mitophagy intermediates from EM analysis of Vps13D depleted neurons. **A)** Two additional examples of larval neurons expressing *Vps13D*-RNAi which contain mitochondria engaged with a phagophore. In the lower magnification images on the left side, mitochondria (M) that are not engaged with a phagophore have compact and electron dense cristae. In comparison, the phagophore-associated mitochondria (M_p_) have cristae that appear less compactly organized. High magnification image of mitophagy intermediates (dashed black box) are shown to the right. Arrowheads indicate the phagophore. **B)** Two additional examples of mitochondrial rupture by phagophore-associated mitochondria (M_p_) in *Vps13D*-RNAi neurons. Arrowheads indicate the phagophore, while arrows indicate locations on the mitochondria lacking IMM and OMM. **C)** Additional example of micrograph from a *Vps13D*-RNAi neuron: the two mitochondria in the same neuron show some differences in cristae organization and electron density.

**Figure 2, Figure Supplement 2:**
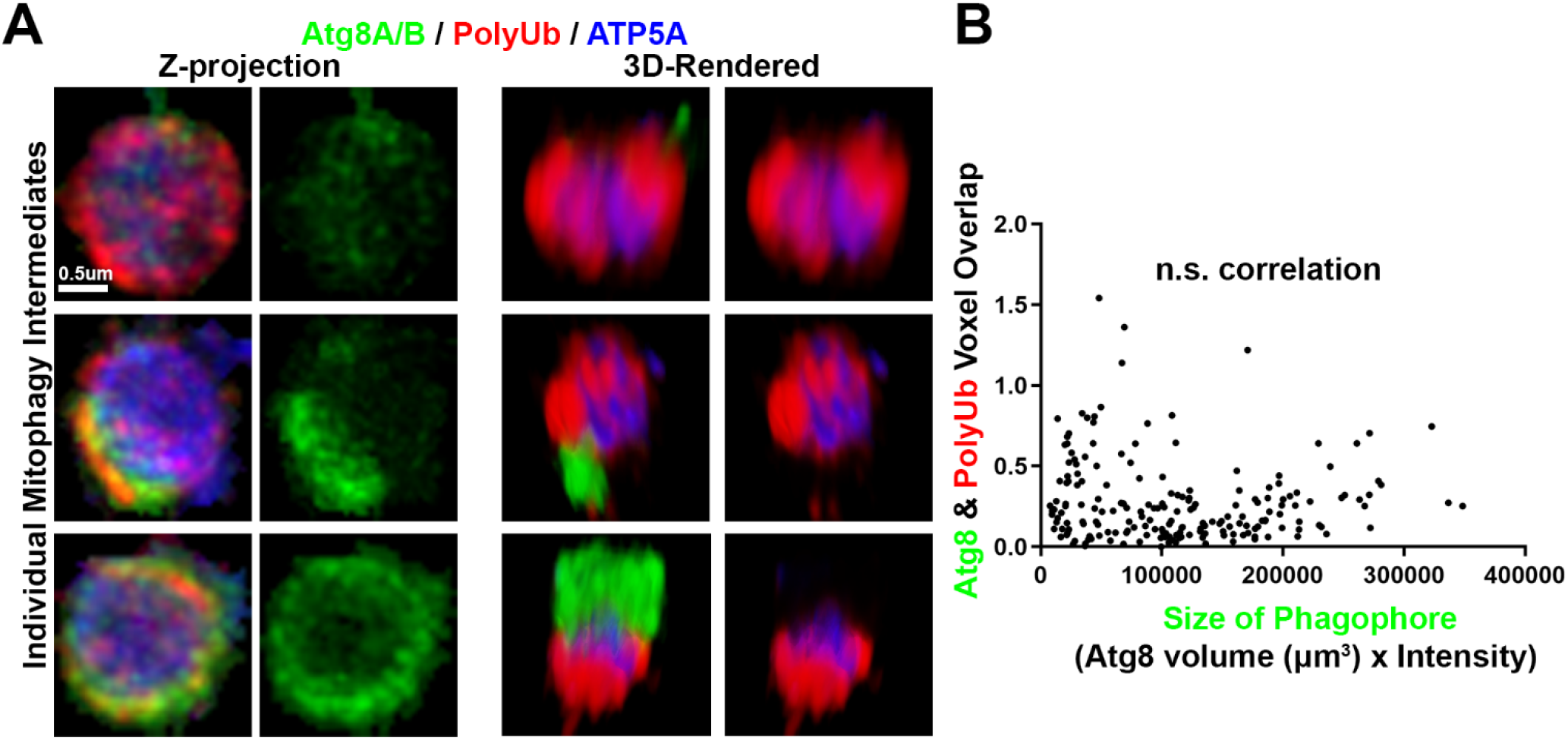
Partial phagophores associated with mitophagy intermediates in Vps13D depleted neurons. **A)** Representative images of individual mitophagy intermediates engaged with various sized phagophores (smallest to largest from top to bottom). Mitophagy intermediates were stained for mitochondrial marker ATP5A (blue), polyubiquitin (PolyUb) (red), and phagophore protein Atg8A/B (green). Left panel shows projected confocal image, and 3D renderings of the projections are shown in the middle and right panels to portray the shape of the engaged phagophore on the mitophagy intermediate. **B)** Quantitative analysis of phagophore engulfment of mitophagy intermediates. The sum of Atg8 staining per mitophagy intermediate (Volume x Intensity) is plotted on the X-axis against the voxel overlap of Atg8 (green) and polyubiquitin (red). Each point represents a single mitophagy intermediate. (n.s. correlation indications non-significance in Pearson’s Correlation test, n=194 XY pairs collected from 5 larvae VNCs, p=0.3). If the phagophore was successfully engulfing the mitochondria, we expect there would be a positive correlation between the size of the phagophore and the overlap with PolyUb.

**Figure 3, Figure Supplement 1:**
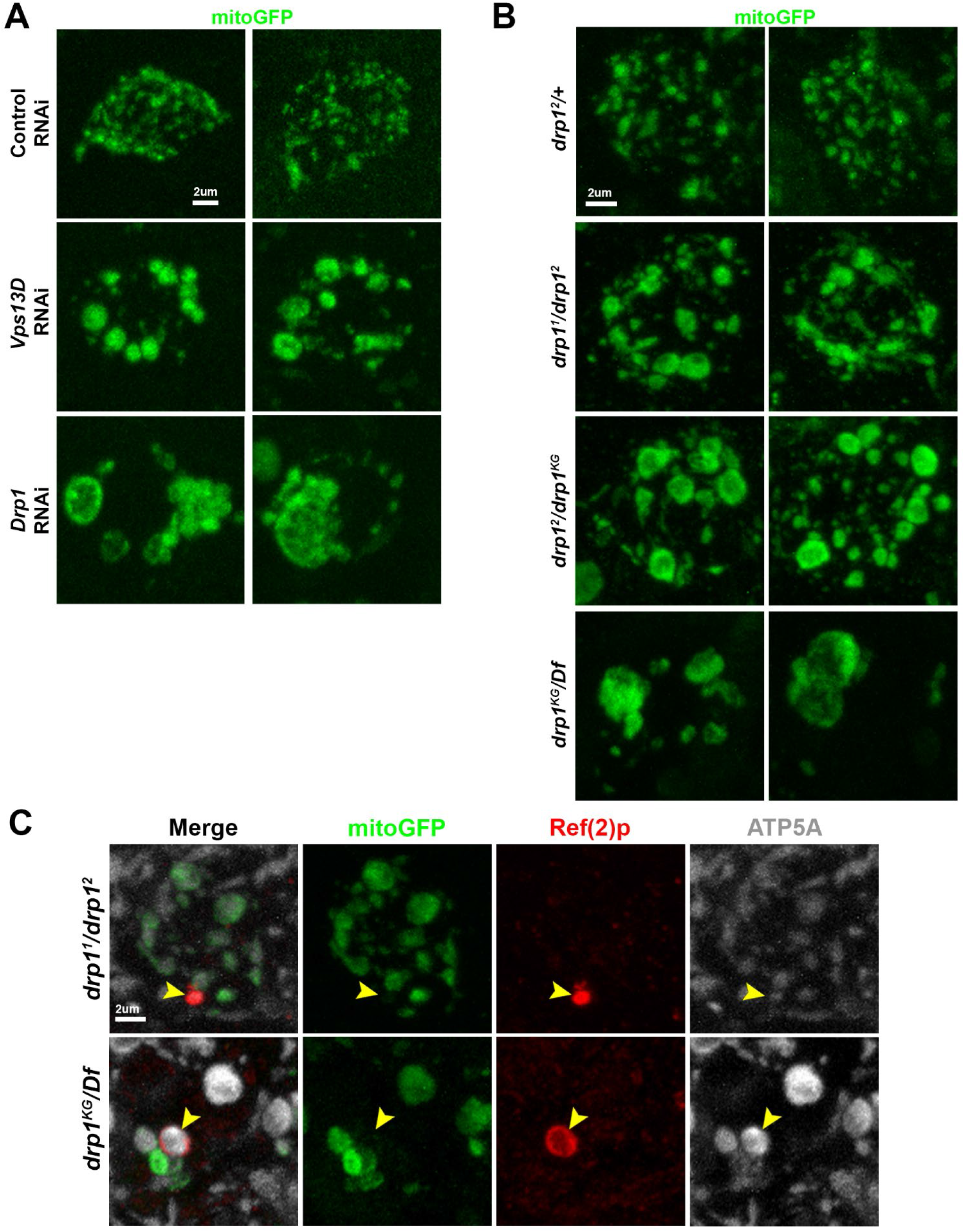
Further characterization of *drp1* mutant motoneurons. **A)** Representative images of individual dorsal midline motoneuron cell bodies which co-express the indicated RNAi together with mitoGFP (green) driven by the pan-motoneuron driver D42-Gal4. RNAi depletion of either Vps13D (BL# 38320) or Drp1 (BL# 67160) leads to enlarged mitochondrial morphology. **B)** Representative images of individual dorsal midline motoneuron cell bodies from the indicated genotypes which express mitoGFP (green) driven by the pan-motoneuron driver D42-Gal4. Enlargement of mitochondrial morphology is most severe and similar to *Drp1* RNAi condition in the *drp1^KG^/Df* genotype (bottom). Morphological enlargement is more pronounced in *drp1^2^/drp1^KG^* compared to *drp1^2^/drp1^2^* genotype. **C)** Representative images of motoneurons in the larval VNC of indicated *drp1* mutant which express mitochondrial marker mitoGFP (green) via the D42-Gal4 driver, which is stained for Ref(2)p (red) and ATP5A (greyscale). Yellow arrowheads indicate example mitophagy intermediates lacking mitochondrial matrix (Ref(2)p+/ATP5A+/mitoGFP-).

**Figure 4, Figure Supplement 1:**
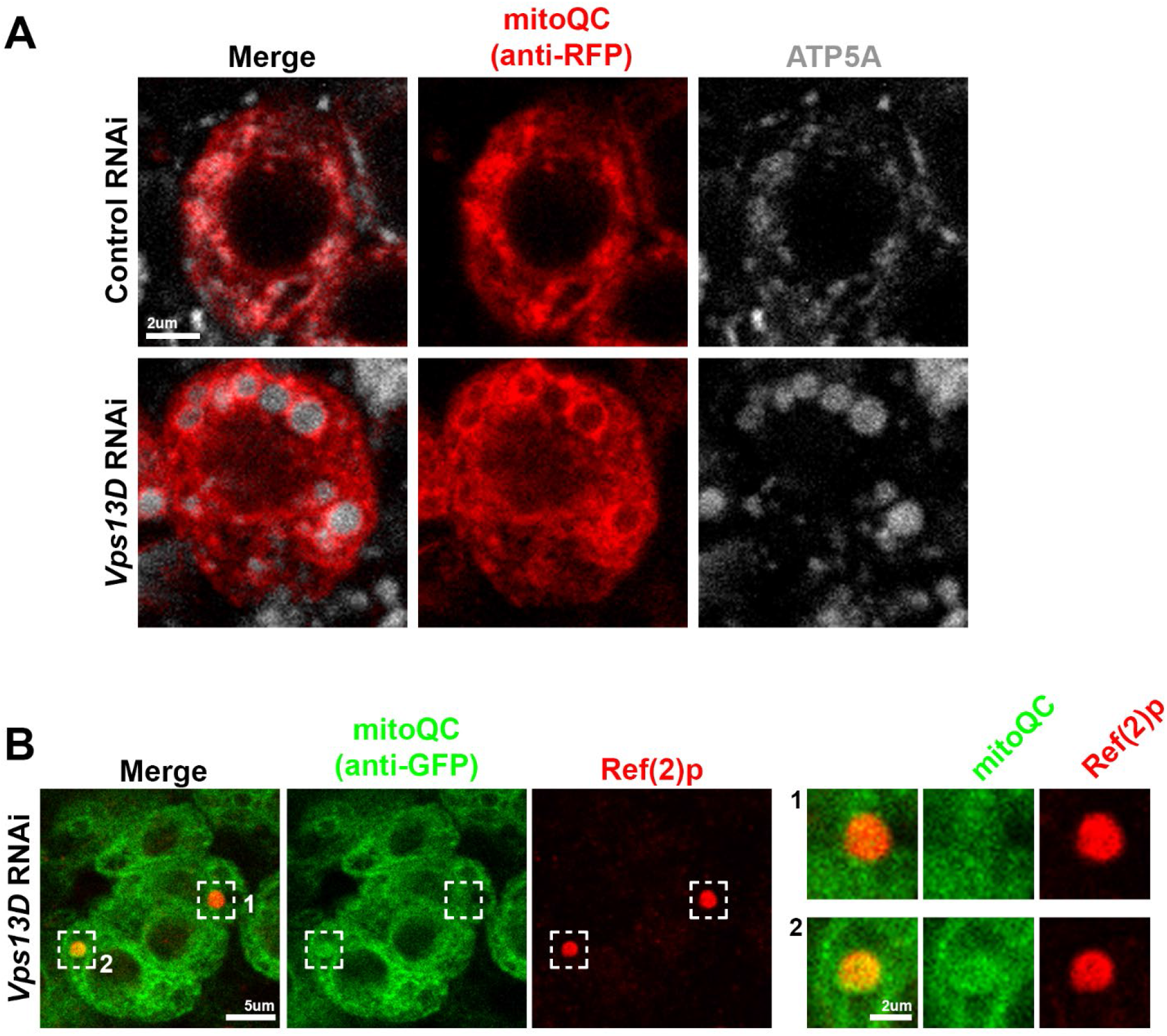
mitoQC marker does not consistently label mitochondria or mitophagy intermediates in larval motoneurons. **A)** Representative images of individual dorsal midline motoneuron cell bodies co-expressing the indicated RNAi and UAS-mitoQC via the D42-Gal4. Tissue was fixed and stained with antibodies against RFP to recognize mitoQC reporter (red) and ATP5A to label the mitochondria. While mitoQC concentrates on mitochondria in both knockdown conditions, it also labels much of the cytoplasm. **B)** Representative images of individual dorsal midline motoneuron cell bodies co-expressing the *Vps13D*-RNAi and UAS-mitoQC via the D42-Gal4. Tissue was fixed and stained with antibodies against GFP to recognize mitoQC reporter (green) and Ref(2) to label the mitophagy intermediates. While mitoQC sometimes concentrated on mitophagy intermediates (example #2), it did not consistently label mitophagy intermediates in Vps13D depleted neurons (example #1).

**Figure 4, Figure Supplement 2:**
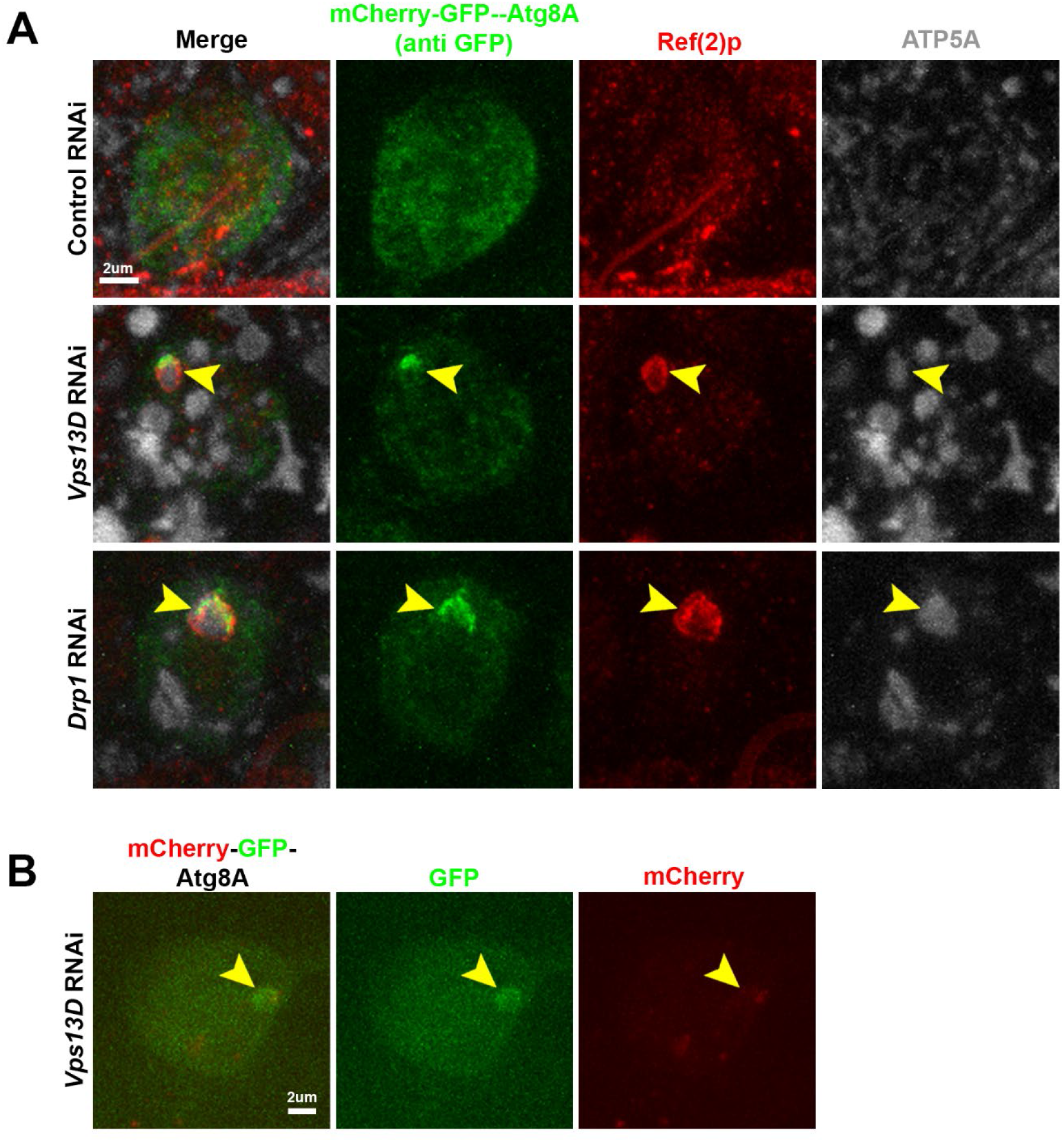
Further analysis of mCherry-GFP-Atg8A autophagy reporter. **A)** Representative images of individual dorsal midline motoneuron cell bodies co-expressing the indicated RNAi and UAS-mCherry-GFP-Atg8A via the D42-Gal4 driver. Tissue was fixed and stained with antibodies against GFP to recognize the tandem-tagged Atg8A reporter (green), Ref(2)p (red), and ATP5A (greyscale) to label the mitochondria. The reporter consistently localizes to mitophagy intermediates (yellow arrowheads) in *Vps13D* and *Drp1* RNAi conditions, consistent with endogenous Atg8A/B staining. **B)** Representative image of live individual dorsal midline motoneuron cell body co-expressing *Vps13D* RNAi and UAS-mCherry-GFP-Atg8A via the D42-Gal4. Yellow arrowhead indicates rounded concentration of reporter, presumably associated with a mitochondrion (though not labeled in this live imaging experiment). While GFP accumulation is obvious, expected RFP accumulation is difficult to observe in this image due to the imaging conditions necessary to capture the significantly brighter red-only puncta without saturating pixels.

**Figure 4, Figure Supplement 3:**
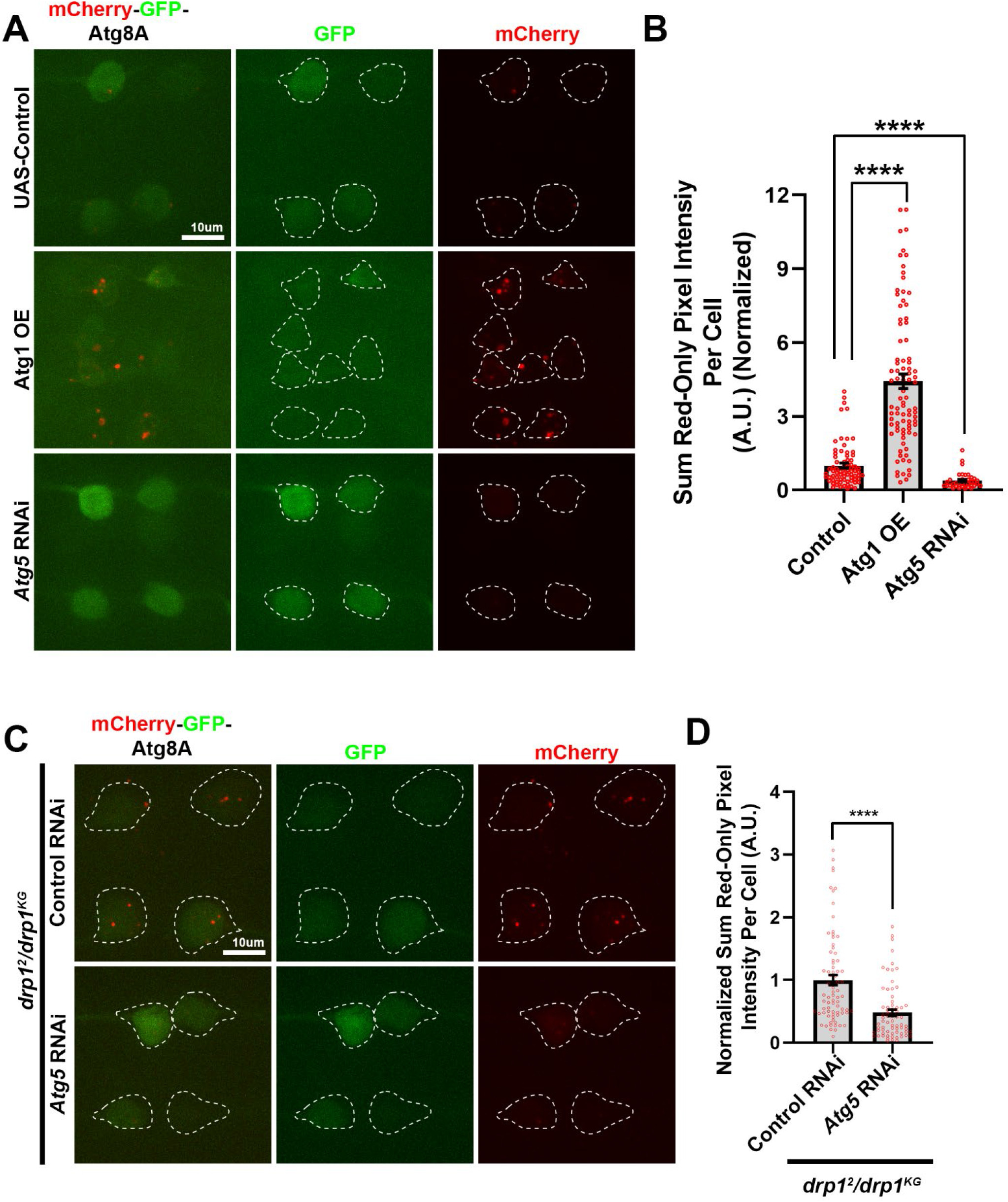
Control conditions to verify tandem-tagged reporter results in larval motoneurons. **A)** Representative images of live larval motoneurons expressing UAS-mCherry-GFP-Atg8A, expressed via the D42-Gal4 driver, simultaneous with expression of the indicated protein overexpression (OE) or RNAi. White dashed lines indicate the outlines of individual cell bodies. **B)** Quantification of the sum pixel intensity of the red-only signal per neuronal cell body (normalized to UAS-Control (UAS-luciferase)). Each point represents a single neuronal cell body, bars represent the mean ± SEM (Control n=74 cell bodies; Atg1 OE n=89 cell bodies; and *Atg5* RNAi n=37 cell bodies from 6 larval VNCs each). **** represents p value <0.0001. **C)** Representative images of live larval motoneurons co-expressing UAS-mCherry-GFP-Atg8A along with indicated RNAi, via the D42-Gal4, in *drp1* mutants (*drp1^2^/drp1^KG^)*. White dashed lines indicate the outlines of individual cell bodies. **D)** Quantification of the sum pixel intensity of the red-only signal per neuronal cell body (normalized to Control RNAi in *drp1* mutant). Each point represents a single neuronal cell body, bars represent the mean ± SEM (Control RNAi n=76 cell bodies; *Atg5* RNAi n=72 cell bodies from 7 larval VNCs each). **** represents p value <0.0001.

**Figure 5, Figure Supplement 1:**
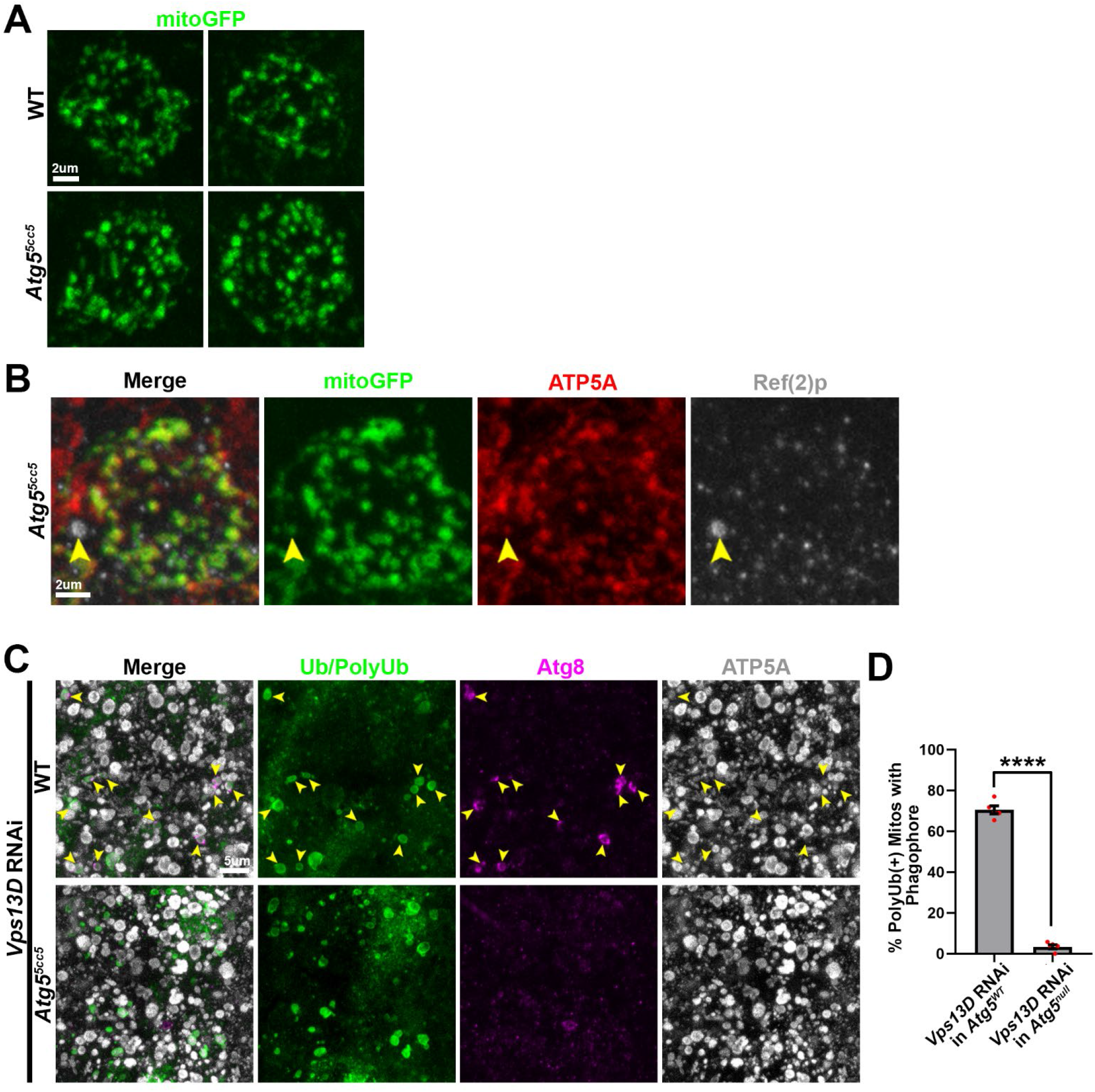
Further analysis and validation of mitophagy blockage in *Atg5* null mutants. **A,B)** Representative images of individual dorsal midline motoneuron cell bodies which express mitoGFP (green) driven by the pan-motoneuron driver D42-Gal4 in WT (*w^1118^*) (top) and *Atg5* null (*Atg5^5cc5^*) animals (bottom). **B)** Tissue was stained for the mitochondrial protein ATP5A (red) and autophagy receptor Ref(2)p (greyscale). Yellow arrowhead indicates a mitophagy intermediate which contains both ATP5A and mitoGFP. **C)** Representative images of dorsal midline motoneurons which express *Vps13D* RNAi driven by pan-neuronal driver Elav-Gal4 in a WT (*w^1118^*) background (top panel) vs. an *Atg5* null (*Atg5^5cc5^*) background (bottom panel). Tissue was stained for ubiquitin (Ub/PolyUb, FK2) (green), phagophore protein Atg8A/B (magenta) and mitochondrial protein ATP5A (greyscale). Yellow arrowheads indicate polyubiquitinated mitochondria engaged with a phagophore (top panel). **D)** Quantification of the % of polyubiquitinated mitochondria that are engaged with a phagophore. Each red point represents the total percentage in the VNC from one animal, and bars represent mean ± SEM. (n=5 for each condition, each containing >50 polyubiquitinated mitochondria). **** indicates p<0.0001.

**Figure 5, Figure Supplement 2:**
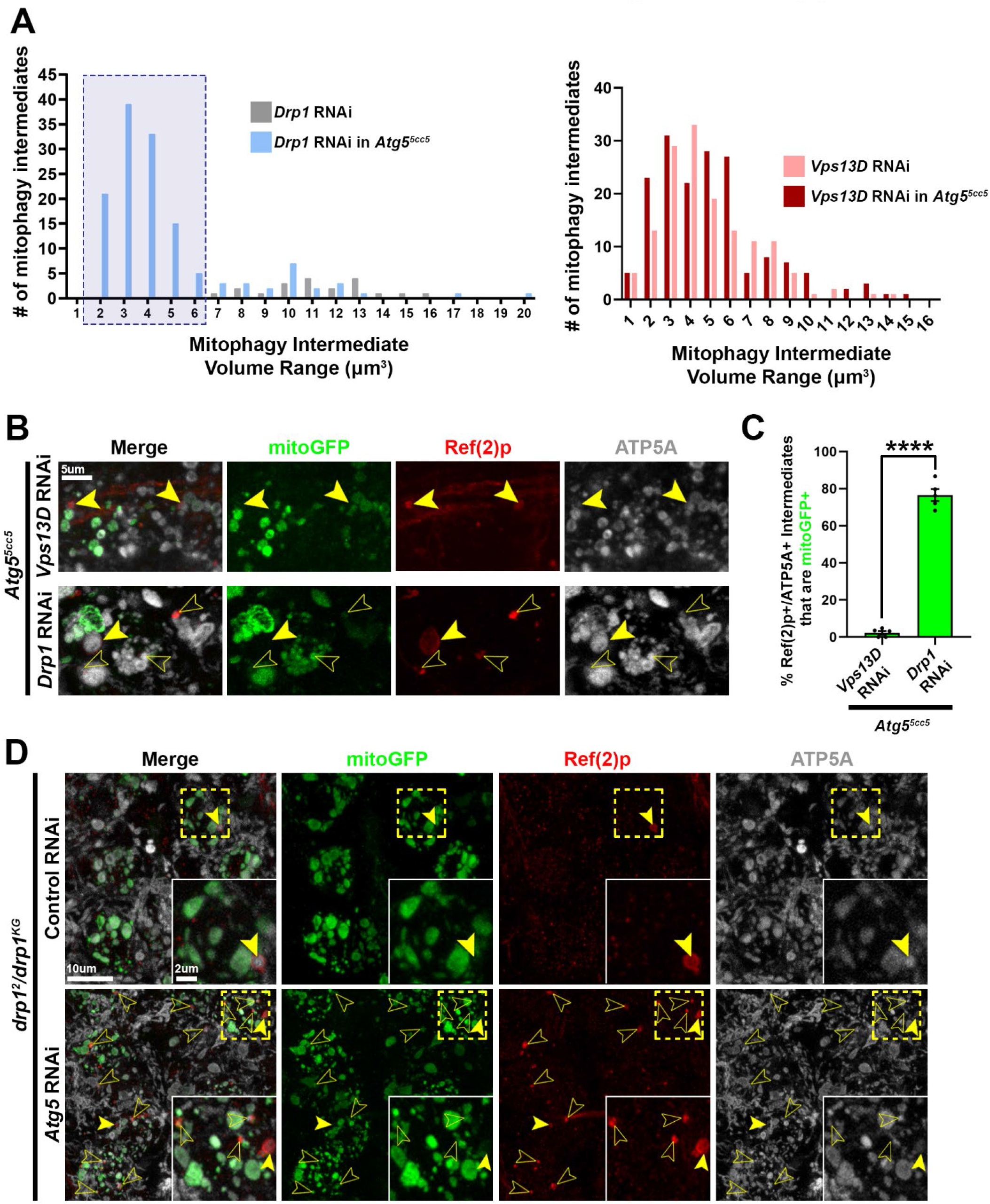
Mitophagy intermediates in conditions of combined Drp1 and Atg5 loss. **A)** Histograms depicting the distribution of the volume (μm^3^) of mitophagy intermediates (Ref(2)p+ mitochondria) in fission-deficient conditions in WT and *Atg5* mutant backgrounds. Top histogram represents conditions of *Drp1* RNAi expression (n=20 mitophagy intermediate for *Drp1* RNAi condition (grey bars), and n=136 mitophagy intermediates for *Drp1* RNAi in *Atg5* mutant condition (light blue bars). Bottom histogram represents *Vps13D* RNAi expression (n=168 mitophagy intermediates for *Vps13D* RNAi condition (pink bars) and n=144 mitophagy intermediates in *Vps13D* RNAi condition in *Atg5* mutants (red bars)). Blue shaded box with dashed lines indicates a population of smaller mitophagy intermediates that are revealed in Drp1 depleted neurons only when *Atg5* is lost. No such new population of mitophagy intermediates is revealed when Vps13D is depleted in *Atg5* mutant conditions. **B)** Representative images of dorsal midline motoneurons from the *Atg5* mutants (*Atg5^5cc5^/y*) which co-express mitoGFP and the indicated RNAi driven by the pan-neuron driver elav-Gal4. Closed yellow arrowheads indicate stalled mitophagy intermediate lacking mitoGFP, whereas open yellow arrowheads indicate stalled mitophagy intermediates containing mitoGFP. **C)** Quantification of the % of stalled mitophagy intermediates that contain mitoGFP. Verified Ref(2)p+/ATP5A+ objects were designated as mitoGFP+ based on described method in **Figure 1C**. Points represent the % of mitoGFP+ mitophagy intermediates out of total Ref(2)p+/ATP5A+ mitophagy intermediates in one animal, with n=5 animals per RNAi condition (each condition contained >160 Ref(2)p+/ATP5A+ mitophagy intermediates). Bars represent mean ± SEM. **** indicates p<0.0001. **D)** Representative images of dorsal midline motoneurons from the *drp1* mutants (*drp1^2^/drp1^KG^*) which co-express mitoGFP and the indicated RNAi driven by the pan-motoneuron driver D42-Gal4. Dashed yellow box outlines a single Gal4-expressing neuronal cell body that is shown in high magnification in the inset in the bottom right corner. Yellow arrowheads indicate example mitophagy intermediates (Ref(2)p+/ATP5A+). In inset of *drp1* mutant expressing *Atg5* RNAi (bottom), open yellow arrowheads indicate mitophagy intermediates that contain mitoGFP, while the closed arrowhead indicates mitophagy intermediates lacking mitoGFP.

## References

Adachi, Yoshihiro, Kie Itoh, Tatsuya Yamada, Kara L. Cerveny, Takamichi L. Suzuki, Patrick Macdonald, Michael A. Frohman, Rajesh Ramachandran, Miho Iijima, and Hiromi Sesaki. 2016. “Coincident Phosphatidic Acid Interaction Restrains Drp1 in Mitochondrial Division.” Molecular Cell 63 (6): 1034–43.

Allen, George F. G., Rachel Toth, John James, and Ian G. Ganley. 2013. “Loss of Iron Triggers PINK1/Parkin-Independent Mitophagy.” EMBO Reports 14 (12): 1127–35.

Anding, Allyson L., Chunxin Wang, Tsun-Kai Chang, Danielle A. Sliter, Christine M. Powers, Kay Hofmann, Richard J. Youle, and Eric H. Baehrecke. 2018. “Vps13D Encodes a Ubiquitin-Binding Protein That Is Required for the Regulation of Mitochondrial Size and Clearance.” Current Biology: CB 28 (2): 287–95.e6.

Ariosa, Aileen R., and Daniel J. Klionsky. 2016. “Autophagy Core Machinery: Overcoming Spatial Barriers in Neurons.” Journal of Molecular Medicine 94 (11): 1217–27.

Barekat, Ayeh, Arysa Gonzalez, Ruth E. Mauntz, Roxanne W. Kotzebue, Brandon Molina, Nadja El-Mecharrafie, Catherine J. Conner, et al. 2016. “Using Drosophila as an Integrated Model to Study Mild Repetitive Traumatic Brain Injury.” Scientific Reports 6 (May): 25252.

Bean, Björn D. M., Samantha K. Dziurdzik, Kathleen L. Kolehmainen, Claire M. S. Fowler, Waldan K. Kwong, Leslie I. Grad, Michael Davey, Cayetana Schluter, and Elizabeth Conibear. 2018. “Competitive Organelle-Specific Adaptors Recruit Vps13 to Membrane Contact Sites.” The Journal of Cell Biology 217 (10): 3593–3607.

Bellen, Hugo J., Robert W. Levis, Guochun Liao, Yuchun He, Joseph W. Carlson, Garson Tsang, Martha Evans-Holm, et al. 2004. “The BDGP Gene Disruption Project: Single Transposon Insertions Associated with 40% of Drosophila Genes.” Genetics 167 (2): 761–81.

Bendotti, C., N. Calvaresi, L. Chiveri, A. Prelle, M. Moggio, M. Braga, V. Silani, and S. De Biasi. 2001. “Early Vacuolization and Mitochondrial Damage in Motor Neurons of FALS Mice Are Not Associated with Apoptosis or with Changes in Cytochrome Oxidase Histochemical Reactivity.” Journal of the Neurological Sciences 191 (1-2): 25–33.

Burman, Jonathon L., Sarah Pickles, Chunxin Wang, Shiori Sekine, Jose Norberto S. Vargas, Zhe Zhang, Alice M. Youle, et al. 2017. “Mitochondrial Fission Facilitates the Selective Mitophagy of Protein Aggregates.” The Journal of Cell Biology 216 (10): 3231–47.

Burté, Florence, Valerio Carelli, Patrick F. Chinnery, and Patrick Yu-Wai-Man. 2015. “Disturbed Mitochondrial Dynamics and Neurodegenerative Disorders.” Nature Reviews. Neurology 11 (1): 11–24.

Cai, Qian, Hesham Mostafa Zakaria, Anthony Simone, and Zu-Hang Sheng. 2012. “Spatial Parkin Translocation and Degradation of Damaged Mitochondria via Mitophagy in Live Cortical Neurons.” Current Biology: CB 22 (6): 545–52.

Cao, Xu, Haiqiong Wang, Zhao Wang, Qingyao Wang, Shuang Zhang, Yuanping Deng, and Yanshan Fang. 2017. “In Vivo Imaging Reveals Mitophagy Independence in the Maintenance of Axonal Mitochondria during Normal Aging.” Aging Cell 16 (5): 1180–90.

Chan, Nickie C., Anna M. Salazar, Anh H. Pham, Michael J. Sweredoski, Natalie J. Kolawa, Robert L. J. Graham, Sonja Hess, and David C. Chan. 2011. “Broad Activation of the Ubiquitin-Proteasome System by Parkin Is Critical for Mitophagy.” Human Molecular Genetics 20 (9): 1726–37.

Cornelissen, Tom, Sven Vilain, Katlijn Vints, Natalia Gounko, Patrik Verstreken, and Wim Vandenberghe. 2018. “Deficiency of Parkin and PINK1 Impairs Age-Dependent Mitophagy in Drosophila.” eLife 7 (May). https://doi.org/10.7554/eLife.35878.

Cummins, Nadia, and Jürgen Götz. 2018. “Shedding Light on Mitophagy in Neurons: What Is the Evidence for PINK1/Parkin Mitophagy in Vivo?” Cellular and Molecular Life Sciences: CMLS 75 (7): 1151–62.

Davies, Vanessa J., Andrew J. Hollins, Malgorzata J. Piechota, Wanfen Yip, Jennifer R. Davies, Kathryn E. White, Phillip P. Nicols, Michael E. Boulton, and Marcela Votruba. 2007. “Opa1 Deficiency in a Mouse Model of Autosomal Dominant Optic Atrophy Impairs Mitochondrial Morphology, Optic Nerve Structure and Visual Function.” Human Molecular Genetics 16 (11): 1307–18.

Detmer, Scott A., Christine Vande Velde, Don W. Cleveland, and David C. Chan. 2008. “Hindlimb Gait Defects due to Motor Axon Loss and Reduced Distal Muscles in a Transgenic Mouse Model of Charcot-Marie-Tooth Type 2A.” Human Molecular Genetics 17 (3): 367–75.

Duncan, Dianne M., Paula Kiefel, and Ian Duncan. 2017. “Mutants for Drosophila Isocitrate Dehydrogenase 3b Are Defective in Mitochondrial Function and Larval Cell Death.” G3 7 (3): 789–99.

El Fissi, Najla, Manuel Rojo, Aїcha Aouane, Esra Karatas, Gabriela Poliacikova, Claudine David, Julien Royet, and Thomas Rival. 2018. “Mitofusin Gain and Loss of Function Drive Pathogenesis in Drosophila Models of CMT2A Neuropathy.” EMBO Reports 19 (8). https://doi.org/10.15252/embr.201745241.

Frank, Magdalena, Stéphane Duvezin-Caubet, Sebastian Koob, Angelo Occhipinti, Ravi Jagasia, Anton Petcherski, Mika O. Ruonala, Muriel Priault, Bénédicte Salin, and Andreas S. Reichert. 2012. “Mitophagy Is Triggered by Mild Oxidative Stress in a Mitochondrial Fission Dependent Manner.” Biochimica et Biophysica Acta (BBA) - Molecular Cell Research 1823 (12): 2297–2310.

Friedman, Jonathan R., Laura L. Lackner, Matthew West, Jared R. DiBenedetto, Jodi Nunnari, and Gia K. Voeltz. 2011. “ER Tubules Mark Sites of Mitochondrial Division.” Science 334 (6054): 358–62.

Gatica, Damián, Vikramjit Lahiri, and Daniel J. Klionsky. 2018. “Cargo Recognition and Degradation by Selective Autophagy.” Nature Cell Biology 20 (3): 233–42.

Gauthier, Julie, Inge A. Meijer, Davor Lessel, Niccolò E. Mencacci, Dimitri Krainc, Maja Hempel, Konstantinos Tsiakas, et al. 2018. “Recessive Mutations in >VPS13D Cause Childhood Onset Movement Disorders.” Annals of Neurology 83 (6): 1089–95.

Geisler, Sven, Kira M. Holmström, Diana Skujat, Fabienne C. Fiesel, Oliver C. Rothfuss, Philipp J. Kahle, and Wolfdieter Springer. 2010. “PINK1/Parkin-Mediated Mitophagy Is Dependent on VDAC1 and p62/SQSTM1.” Nature Cell Biology 12 (2): 119–31.

Higgins, Cynthia M. J., Cheolwha Jung, and Zuoshang Xu. 2003. “ALS-Associated Mutant SOD1G93A Causes Mitochondrial Vacuolation by Expansion of the Intermembrane Space and by Involvement of SOD1 Aggregation and Peroxisomes.” BMC Neuroscience 4 (July): 16.

Kageyama, Yusuke, Masahiko Hoshijima, Kinya Seo, Djahida Bedja, Polina Sysa-Shah, Shaida A. Andrabi, Weiran Chen, et al. 2014. “Parkin-Independent Mitophagy Requires Drp1 and Maintains the Integrity of Mammalian Heart and Brain.” The EMBO Journal 33 (23): 2798–2813.

Kageyama, Yusuke, Zhongyan Zhang, Ricardo Roda, Masahiro Fukaya, Junko Wakabayashi, Nobunao Wakabayashi, Thomas W. Kensler, P. Hemachandra Reddy, Miho Iijima, and Hiromi Sesaki. 2012. “Mitochondrial Division Ensures the Survival of Postmitotic Neurons by Suppressing Oxidative Damage.” The Journal of Cell Biology 197 (4): 535–51.

Katayama, Hiroyuki, Hiroshi Hama, Koji Nagasawa, Hiroshi Kurokawa, Mayu Sugiyama, Ryoko Ando, Masaaki Funata, et al. 2020. “Visualizing and Modulating Mitophagy for Therapeutic Studies of Neurodegeneration.” Cell 181 (5): 1176–87.e16.

Kim, Myungjin, Erin Sandford, Damian Gatica, Yu Qiu, Xu Liu, Yumei Zheng, Brenda A. Schulman, et al. 2016. “Mutation in ATG5 Reduces Autophagy and Leads to Ataxia with Developmental Delay.” eLife 5 (January). https://doi.org/10.7554/eLife.12245.

Kimura, Shunsuke, Takeshi Noda, and Tamotsu Yoshimori. 2007. “Dissection of the Autophagosome Maturation Process by a Novel Reporter Protein, Tandem Fluorescent-Tagged LC3.” Autophagy 3 (5): 452–60.

Kinnally, Kathleen W., Pablo M. Peixoto, Shin-Young Ryu, and Laurent M. Dejean. 2011. “Is mPTP the Gatekeeper for Necrosis, Apoptosis, or Both?” Biochimica et Biophysica Acta 1813 (4): 616–22.

Klionsky, Daniel J., Kotb Abdelmohsen, Akihisa Abe, Md Joynal Abedin, Hagai Abeliovich, Abraham Acevedo Arozena, Hiroaki Adachi, et al. 2016. “Guidelines for the Use and Interpretation of Assays for Monitoring Autophagy (3rd Edition).” Autophagy 12 (1): 1–222.

Koh, Kishin, Hiroyuki Ishiura, Haruo Shimazaki, Michiko Tsutsumiuchi, Yuta Ichinose, Haitian Nan, Shun Hamada, Toshihisa Ohtsuka, Shoji Tsuji, and Yoshihisa Takiyama. 2020. “VPS13D-Related Disorders Presenting as a Pure and Complicated Form of Hereditary Spastic Paraplegia.” Molecular Genetics & Genomic Medicine 8 (3): e1108.

Kong, J., and Z. Xu. 1998. “Massive Mitochondrial Degeneration in Motor Neurons Triggers the Onset of Amyotrophic Lateral Sclerosis in Mice Expressing a Mutant SOD1.” The Journal of Neuroscience: The Official Journal of the Society for Neuroscience 18 (9): 3241–50.

Kumar, Nikit, Marianna Leonzino, William Hancock-Cerutti, Florian A. Horenkamp, Peiqi Li, Joshua A. Lees, Heather Wheeler, Karin M. Reinisch, and Pietro De Camilli. 2018. “VPS13A and VPS13C Are Lipid Transport Proteins Differentially Localized at ER Contact Sites.” The Journal of Cell Biology 217 (10): 3625–39.

Lee, Juliette J., Alvaro Sanchez-Martinez, Aitor Martinez Zarate, Cristiane Benincá, Ugo Mayor, Michael J. Clague, and Alexander J. Whitworth. 2018. “Basal Mitophagy Is Widespread in Drosophila but Minimally Affected by Loss of Pink1 or Parkin.” The Journal of Cell Biology 217 (5): 1613–22.

Lesage, Suzanne, Valérie Drouet, Elisa Majounie, Vincent Deramecourt, Maxime Jacoupy, Aude Nicolas, Florence Cormier-Dequaire, et al. 2016. “Loss of VPS13C Function in Autosomal-Recessive Parkinsonism Causes Mitochondrial Dysfunction and Increases PINK1/Parkin-Dependent Mitophagy.” American Journal of Human Genetics 98 (3): 500–513.

Li, Jiaxing, Yao V. Zhang, Elham Asghari Adib, Doychin T. Stanchev, Xin Xiong, Susan Klinedinst, Pushpanjali Soppina, et al. 2017. “Restraint of Presynaptic Protein Levels by Wnd/DLK Signaling Mediates Synaptic Defects Associated with the Kinesin-3 Motor Unc-104.” eLife 6 (September). https://doi.org/10.7554/eLife.24271.

Li, Peiqi, Joshua Aaron Lees, C. Patrick Lusk, and Karin M. Reinisch. 2020. “Cryo-EM Reconstruction of a VPS13 Fragment Reveals a Long Groove to Channel Lipids between Membranes.” The Journal of Cell Biology 219 (5). https://doi.org/10.1083/jcb.202001161.

Lu, Qun, Peiguo Yang, Xinxin Huang, Wanqiu Hu, Bin Guo, Fan Wu, Long Lin, Attila L. Kovács, Li Yu, and Hong Zhang. 2011. “The WD40 Repeat PtdIns(3)P-Binding Protein EPG-6 Regulates Progression of Omegasomes to Autophagosomes.” Developmental Cell 21 (2): 343–57.

MacVicar, Thomas D. B., and Jon D. Lane. 2014. “Impaired OMA1-Dependent Cleavage of OPA1 and Reduced DRP1 Fission Activity Combine to Prevent Mitophagy in Cells That Are Dependent on Oxidative Phosphorylation.” Journal of Cell Science 127 (10): 2313–25.

Markaki, Maria, and Nektarios Tavernarakis. 2020. “Mitochondrial Turnover and Homeostasis in Ageing and Neurodegeneration.” FEBS Letters, April. https://doi.org/10.1002/1873-3468.13802.

McWilliams, Thomas G., Alan R. Prescott, George F. G. Allen, Jevgenia Tamjar, Michael J. Munson, Calum Thomson, Miratul M. K. Muqit, and Ian G. Ganley. 2016. “Mito-QC Illuminates Mitophagy and Mitochondrial Architecture in Vivo.” The Journal of Cell Biology 214 (3): 333–45.

McWilliams, Thomas G., Alan R. Prescott, Lambert Montava-Garriga, Graeme Ball, François Singh, Erica Barini, Miratul M. K. Muqit, Simon P. Brooks, and Ian G. Ganley. 2018. “Basal Mitophagy Occurs Independently of PINK1 in Mouse Tissues of High Metabolic Demand.” Cell Metabolism 27 (2): 439–49.e5.

Misgeld, Thomas, and Thomas L. Schwarz. 2017. “Mitostasis in Neurons: Maintaining Mitochondria in an Extended Cellular Architecture.” Neuron 96 (3): 651–66.

Muñoz-Braceras, Sandra, Rosa Calvo, and Ricardo Escalante. 2015. “TipC and the Chorea-Acanthocytosis Protein VPS13A Regulate Autophagy in Dictyostelium and Human HeLa Cells.” Autophagy 11 (6): 918–27.

Nagy, Péter, Ágnes Varga, Attila L. Kovács, Szabolcs Takáts, and Gábor Juhász. 2015. “How and Why to Study Autophagy in Drosophila: It’s More than Just a Garbage Chute.” Methods 75 (March): 151–61.

Narendra, Derek P., Seok Min Jin, Atsushi Tanaka, Der-Fen Suen, Clement A. Gautier, Jie Shen, Mark R. Cookson, and Richard J. Youle. 2010. “PINK1 Is Selectively Stabilized on Impaired Mitochondria to Activate Parkin.” PLoS Biology 8 (1): e1000298.

Narendra, Derek, Atsushi Tanaka, Der-Fen Suen, and Richard J. Youle. 2008. “Parkin Is Recruited Selectively to Impaired Mitochondria and Promotes Their Autophagy.” The Journal of Cell Biology 183 (5): 795–803.

Oettinghaus, B., J. M. Schulz, L. M. Restelli, M. Licci, C. Savoia, A. Schmidt, K. Schmitt, et al. 2016. “Synaptic Dysfunction, Memory Deficits and Hippocampal Atrophy due to Ablation of Mitochondrial Fission in Adult Forebrain Neurons.” Cell Death and Differentiation 23 (1): 18–28.

Park, Jae-Sook, Mary K. Thorsness, Robert Policastro, Luke L. McGoldrick, Nancy M. Hollingsworth, Peter E. Thorsness, and Aaron M. Neiman. 2016. “Yeast Vps13 Promotes Mitochondrial Function and Is Localized at Membrane Contact Sites.” Molecular Biology of the Cell 27 (15): 2435–49.

Picca, Anna, Riccardo Calvani, Hélio José Coelho-Junior, Francesco Landi, Roberto Bernabei, and Emanuele Marzetti. 2020. “Mitochondrial Dysfunction, Oxidative Stress, and Neuroinflammation: Intertwined Roads to Neurodegeneration.” Antioxidants (Basel, Switzerland) 9 (8). https://doi.org/10.3390/antiox9080647.

Pickles, Sarah, Pierre Vigié, and Richard J. Youle. 2018. “Mitophagy and Quality Control Mechanisms in Mitochondrial Maintenance.” Current Biology: CB 28 (4): R170–85.

Pickrell, Alicia M., and Richard J. Youle. 2015. “The Roles of PINK1, Parkin, and Mitochondrial Fidelity in Parkinson’s Disease.” Neuron 85 (2): 257–73.

Pilling, Aaron D., Dai Horiuchi, Curtis M. Lively, and William M. Saxton. 2006. “Kinesin-1 and Dynein Are the Primary Motors for Fast Transport of Mitochondria in Drosophila Motor Axons.” Molecular Biology of the Cell 17 (4): 2057–68.

Prinz, William A., and James H. Hurley. 2020. “A Firehose for Phospholipids.” The Journal of Cell Biology 219 (5). https://doi.org/10.1083/jcb.202003132.

Rambold, Angelika S., Brenda Kostelecky, Natalie Elia, and Jennifer Lippincott-Schwartz. 2011. “Tubular Network Formation Protects Mitochondria from Autophagosomal Degradation during Nutrient Starvation.” Proceedings of the National Academy of Sciences of the United States of America 108 (25): 10190–95.

Rizzuto, R., M. Brini, P. Pizzo, M. Murgia, and T. Pozzan. 1995. “Chimeric Green Fluorescent Protein as a Tool for Visualizing Subcellular Organelles in Living Cells.” Current Biology: CB 5 (6): 635–42.

Rodger, Catherine E., Thomas G. McWilliams, and Ian G. Ganley. 2018. “Mammalian Mitophagy - from in Vitro Molecules to in Vivo Models.” The FEBS Journal 285 (7): 1185–1202.

Rzepnikowska, Weronika, Krzysztof Flis, Sandra Muñoz-Braceras, Regina Menezes, Ricardo Escalante, and Teresa Zoladek. 2017. “Yeast and Other Lower Eukaryotic Organisms for Studies of Vps13 Proteins in Health and Disease.” Traffic 18 (11): 711–19.

Sandoval, Hector, Chi-Kuang Yao, Kuchuan Chen, Manish Jaiswal, Taraka Donti, Yong Qi Lin, Vafa Bayat, et al. 2014. “Mitochondrial Fusion but Not Fission Regulates Larval Growth and Synaptic Development through Steroid Hormone Production.” eLife 3 (October). https://doi.org/10.7554/eLife.03558.

Seong, Eunju, Ryan Insolera, Marija Dulovic, Erik-Jan Kamsteeg, Joanne Trinh, Norbert Brüggemann, Erin Sandford, et al. 2018. “Mutations in VPS13D Lead to a New Recessive Ataxia with Spasticity and Mitochondrial Defects.” Annals of Neurology, March. https://doi.org/10.1002/ana.25220.

Stewart, B. A., H. L. Atwood, J. J. Renger, J. Wang, and C. F. Wu. 1994. “Improved Stability of Drosophila Larval Neuromuscular Preparations in Haemolymph-like Physiological Solutions A Neuroethology, Sensory, Neural, and Behavioral Physiology.” https://pubag.nal.usda.gov/catalog/1425499.

Sun, Nuo, Jeanho Yun, Jie Liu, Daniela Malide, Chengyu Liu, Ilsa I. Rovira, Kira M. Holmström, et al. 2015. “Measuring In Vivo Mitophagy.” Molecular Cell 60 (4): 685–96.

Takáts, Szabolcs, Péter Nagy, Ágnes Varga, Karolina Pircs, Manuéla Kárpáti, Kata Varga, Attila L. Kovács, Krisztina Hegedűs, and Gábor Juhász. 2013. “Autophagosomal Syntaxin17-Dependent Lysosomal Degradation Maintains Neuronal Function in Drosophila.” The Journal of Cell Biology 201 (4): 531–39.

Tanaka, Atsushi, Megan M. Cleland, Shan Xu, Derek P. Narendra, Der-Fen Suen, Mariusz Karbowski, and Richard J. Youle. 2010. “Proteasome and p97 Mediate Mitophagy and Degradation of Mitofusins Induced by Parkin.” The Journal of Cell Biology 191 (7): 1367–80.

Twig, Gilad, Alvaro Elorza, Anthony J. A. Molina, Hibo Mohamed, Jakob D. Wikstrom, Gil Walzer, Linsey Stiles, et al. 2008. “Fission and Selective Fusion Govern Mitochondrial Segregation and Elimination by Autophagy.” The EMBO Journal 27 (2): 433–46.

Velikkakath, Anoop Kumar G., Taki Nishimura, Eiko Oita, Naotada Ishihara, and Noboru Mizushima. 2012. “Mammalian Atg2 Proteins Are Essential for Autophagosome Formation and Important for Regulation of Size and Distribution of Lipid Droplets.” Molecular Biology of the Cell 23 (5): 896–909.

Verstreken, Patrik, Cindy V. Ly, Koen J. T. Venken, Tong-Wey Koh, Yi Zhou, and Hugo J. Bellen. 2005. “Synaptic Mitochondria Are Critical for Mobilization of Reserve Pool Vesicles at Drosophila Neuromuscular Junctions.” Neuron 47 (3): 365–78.

Vonk, Jan J., Wondwossen M. Yeshaw, Francesco Pinto, Anita I. E. Faber, Liza L. Lahaye, Bart Kanon, Marianne van der Zwaag, et al. 2017. “Drosophila Vps13 Is Required for Protein Homeostasis in the Brain.” PloS One 12 (1): e0170106.

Wei, Yongjie, Wei-Chung Chiang, Rhea Sumpter Jr, Prashant Mishra, and Beth Levine. 2017. “Prohibitin 2 Is an Inner Mitochondrial Membrane Mitophagy Receptor.” Cell 168 (1-2): 224–38.e10.

West, A. Phillip. 2017. “Mitochondrial Dysfunction as a Trigger of Innate Immune Responses and Inflammation.” Toxicology 391 (November): 54–63.

Whitworth, Alexander J., and Leo J. Pallanck. 2017. “PINK1/Parkin Mitophagy and Neurodegeneration-What Do We Really Know in Vivo?” Current Opinion in Genetics & Development 44 (June): 47–53.

Xiao, Bin, Jian-Yuan Goh, Lin Xiao, Hongxu Xian, Kah-Leong Lim, and Yih-Cherng Liou. 2017. “Reactive Oxygen Species Trigger Parkin/PINK1 Pathway-Dependent Mitophagy by Inducing Mitochondrial Recruitment of Parkin.” The Journal of Biological Chemistry 292 (40): 16697–708.

Yamada, Tatsuya, Ted M. Dawson, Toru Yanagawa, Miho Iijima, and Hiromi Sesaki. 2019. “SQSTM1/p62 Promotes Mitochondrial Ubiquitination Independently of PINK1 and PRKN/parkin in Mitophagy.” Autophagy 15 (11): 2012–18.

Yamashita, Shun-Ichi, Xiulian Jin, Kentaro Furukawa, Maho Hamasaki, Akiko Nezu, Hidenori Otera, Tetsu Saigusa, et al. 2016. “Mitochondrial Division Occurs Concurrently with Autophagosome Formation but Independently of Drp1 during Mitophagy.” The Journal of Cell Biology 215 (5): 649–65.

Yeshaw, Wondwossen M., Marianne van der Zwaag, Francesco Pinto, Liza L. Lahaye, Anita Ie Faber, Rubén Gómez-Sánchez, Amalia M. Dolga, et al. 2019. “Human VPS13A Is Associated with Multiple Organelles and Influences Mitochondrial Morphology and Lipid Droplet Motility.” eLife 8 (February). https://doi.org/10.7554/eLife.43561.

Yoshii, Saori R., Chieko Kishi, Naotada Ishihara, and Noboru Mizushima. 2011. “Parkin Mediates Proteasome-Dependent Protein Degradation and Rupture of the Outer Mitochondrial Membrane.” The Journal of Biological Chemistry 286 (22): 19630–40.

Yu, Chien-Hsiung, Sophia Davidson, Cassandra R. Harapas, James B. Hilton, Michael J. Mlodzianoski, Pawat Laohamonthonkul, Cynthia Louis, et al. 2020. “TDP-43 Triggers Mitochondrial DNA Release via mPTP to Activate cGAS/STING in ALS.” Cell 183 (3): 636–49.e18.

Zaninello, Marta, Konstantinos Palikaras, Deborah Naon, Keiko Iwata, Stephanie Herkenne, Ruben Quintana-Cabrera, Martina Semenzato, et al. 2020. “Inhibition of Autophagy Curtails Visual Loss in a Model of Autosomal Dominant Optic Atrophy.” Nature Communications 11 (1): 4029.

